# Intranasal Replicating Adenovirus type 4-SARS-CoV-2 Recombinants Induce Superior Immune Response Durability and Efficacy in Preclinical Testing Compared to Standard Intramuscular Vaccines

**DOI:** 10.64898/2026.02.04.703921

**Authors:** Breanna Kim, Rachel Roenicke, Peyton M. Roeder, Isabel Steinberg, Andy Patamawenu, Mitra Harrison, Freya van’t Veer, Emma Koory, Jane Xie, Tulley Shofner, Kenta Matsuda, Eleanor Wettstein, Ellison Ober, Ceanna Cooney, Tae Kim, Jonathan D. Webber, Tyler Meeks, Tracey Burdette, Shalamar G. Clark, Alexander C. Vostal, Anna Hostal, Jeffrey I. Cohen, Matthew Gagne, Zohar Ziff, Ketan Richard, Matthew R. Burnett, Erin Maule, Viviane Callier, Jason Liang, Gabriela Kovacikova, Scarlett C. Souter, Joshua A. Weiner, Daniel Douek, Margaret E. Ackerman, Peter F. Wright, Reed F. Johnson, Mark Connors

## Abstract

The portfolio of next generation COVID-19 vaccines would benefit from candidates that induce durable systemic and mucosal immune responses that would lessen person-to-person transmission. We constructed an intranasal (IN) replication-competent adenovirus type 4 recombinant platform to express SARS-CoV-2 Spike variants (Ad4-S) and assessed immunogenicity and efficacy in the Syrian hamster model. Although both IN Ad4-S and intramuscular (IM) vaccines (Ad26.CoV2.S and mRNA-1273) induced serum binding antibodies, only Ad4-S induced a robust mucosal response in the nasal cavity. IN Ad4-S vaccination induced serum neutralizing titers equivalent to or greater than IM vaccination but more durable up to 6 months. Upon challenge, IN immunization also resulted in less weight loss, greater breadth and durability of restriction of viral replication, and less lung pathology than IM immunization up to 268 days after immunization. These data support the potential of the IN Ad4 vaccine platform to reduce transmission of SARS-CoV-2 and other respiratory viruses with pandemic potential.

## Introduction

The SARS-CoV-2 pandemic is now in its 6^th^ year and continues to cause serious illness worldwide. Several factors contribute to the persistence of SARS-CoV-2 in the population. Among the most important of these is the development of new variants that are sufficiently genetically distant from prior isolates that they can infect humans with vaccine- or infection-induced immunity. The cycle of infection is perpetuated by the lack of effective means to interrupt transmission, allowing the virus to persist at high levels in the population, and in turn permitting the selection of additional new variants. Another factor is the poor durability of the humoral response induced by some current vaccines. While the durability of vaccine efficacy can be limited by the ongoing development of variants, the situation is further worsened by waning vaccine-induced humoral immunity observed with some current vaccines. For example, in individuals immunized with mRNA vaccines, serum autologous strain-neutralizing antibodies wane substantially by 6 months and protective efficacy against infection following bivalent booster vaccine wanes to 0% by 16 weeks^1^. Thus, development of new vaccines that provide durable immunity and limit new infections and transmission could curtail the development of new strains. While the current portfolio of SARS-CoV-2 vaccines continues to provide protection against severe disease and hospitalization, there are limitations to effectiveness that suggest that additional platforms that might be useful for SARS-CoV-2 or other respiratory pathogens with pandemic potential should be explored.

Among the approaches that might interrupt transmission are those that stimulate local immunity in the respiratory tract through mucosal immunization. In addition to IgG and resident memory T cells, mucosal immunity in the upper respiratory tract can be mediated by dimeric secretory IgA, which can have potent neutralizing capacity^2^. In the nasal cavity, secretory IgA is dependent upon local IgA-producing B cells primarily in the nasal-associated lymphoid tissue^3-5^. Mucosal immunity operates as a compartment separate from systemic immunity and is most potently established by local immunization, although it can be boosted by IM vaccination^6,7^. The SARS-CoV-2 vaccines currently in widespread use, such as mRNA and most replication-defective adenovirus-based vaccines, are administered IM and induce only low mucosal secretory IgA immune responses^8,9^. IM vaccines have induced only low level or inconsistent protection of the upper respiratory tract in experimental animals or humans^10-16^. Conversely, there have been many pre-clinical studies in small animals and Rhesus macaques where mucosally delivered SARS-CoV-2 vaccines induce more potent protection from infection than the same vaccine given IM, macaques^10-16^. In addition, there are several IN or inhaled aerosol platforms that have shown some promise in humans^12,14,17-21^ Although IM SARS-CoV-2 vaccination of humans gives little to no protection from infection, there are some data suggesting that infection induces potent protection from re-infection^22,23^. Taken together the available data strongly suggest that upper respiratory tract immunization may be a powerful tool for limiting infection and transmission.

Over the past decade, our laboratory has developed an IN replication-competant adenovirus type 4 (Ad4) platform to induce systemic and respiratory mucosal immunity. The Ad4 vector used in this approach is based upon the vaccine that has been used in the United States Military for many years with well over 10 million doses administered. In that experience, the wild-type Ad4 is administered via an enteric coated capsule with a remarkably good safety record and 99.3% efficacy^24^. Ad4 vectors expressing influenza virus H5, or HIV genes have now been in six Phase 1 clinical trials, and have proven to be a safe and highly immunogenic. Although the transgene-bearing recombinant vaccine is overly attenuated when administered orally, it remains highly immunogenic after inoculation of the upper respiratory tract. After IN vaccination with Ad4 expressing influenza H5 Vietnam (Ad4-H5), participants developed high levels of serum neutralizing antibodies that were more durable than those induced by standard IM delivered protein vaccines^25-27^. In addition, maturation of the B cell response continued for up to 12 months after a single IN vaccination. The vaccine induced mucosal immunity and the serum neutralizing antibodies were unchanged from their peak after 3-5 years^25^. The Ad4 seroprevalence in the U.S. adults is approximately 30%, which is below that of Ad5, and in some populations Ad26. High dose mucosal administration is a known method to bypass pre-existing systemic immunity^28-30^. Consistent with oral administration of Ad4 in the military, seropositive persons are reinfected by IN administration^25,31^. Given this experience, the platform has highly desirable characteristics as a potential mucosal vaccine for SARS-CoV-2.

In this study we examine the immunogenicity and protective efficacy of IN Ad4 expressing SARS-CoV-2 Spike variants in comparison with the IM Ad26 and mRNA vaccine platforms in hamsters. Unlike IM administered vaccines, the IN Ad4 recombinants reliably induced mucosal antibody responses. In addition, they induced systemic neutralizing antibodies with the same peak level as mRNA but with greater durability at 6 months. Following challenge, animals immunized with Ad4 recombinants had less weight loss, less viral replication, lower levels of pulmonary SARS-CoV-2 antigen and pathology compared to those that received IM vaccines. This protection was quite durable and reduced viral replication and weight loss even 268 days after vaccination in a transmission study. These data, taken in the context of prior clinical experience with other viral transgenes, suggest that IN Ad4 has great promise as a mucosal vaccine for SARS-CoV-2 that may offer both a durable systemic immune response and a protective mucosal immune response that could also reduce transmission.

## Results

The Syrian hamster is an attractive model for investigating mucosal vaccines given they are permissive to SARS-CoV-2 infection at a low inoculum and can readily transmit infection^32^. Although adenovirus replication is typically species-specific, human adenovirus type 5 is known to replicate in Syrian hamsters ^33^. Further, given recent findings that hamsters can be successfully infected intranasally with a replication-defective ChAdOx, that is based upon a Y25 chimpanzee adenovirus that is highly related to Ad4^32^, we sought to investigate whether Ad4 might also replicate, disseminate, and express a transgene in Syrian hamsters. In a dose titration experiment, hamsters were inoculated with a total dose of Ad4-Wuhan ranging from 10^2^-10^7^ infection forming units (IFU). When infected with a total dose of 10^6^ IFU or higher, hamsters demonstrated peak viral DNA in the oropharynx 3 days post-inoculation, that was no longer detected by day 7. Viral replication was not detected at doses of 10^5^ IFU or below. Moreover, the highest levels of Ad4 viral DNA were observed after inoculation with 10^7^ IFU with a mean peak viral load of 1.9 x 10^5^ viral copies/mL 3 days post-infection (Fig. 1A). Only the median area under the curve (AUC) following 10^7^ IFU dose was significantly different from the 10^2^ IFU dose (p=0.041). To determine the duration of expresssion of a transgene, hamsters were intranasally inoculated with 6.8 x 10^6^ IFU of a luciferase-expressing Ad4 and bioluminescence was measured over the next 14 days. Luciferase expression in the nasal cavity was detected from 3 to 5 days post-infection, and decreased to levels that were low or below detection by 14 days post-infection (Fig. 1B, C). We did not observe weight loss following IN inoculation of hamsters with 10^7^ IFU of Ad4 expressing numerous variants of concern (Supplementary Fig. 1). Taken together, the Syrian hamster model is modestly permissive to Ad4 infection in the upper respiratory tract when the inoculum meets or exceeds 10^6^ IFU.

**Figure 1.**
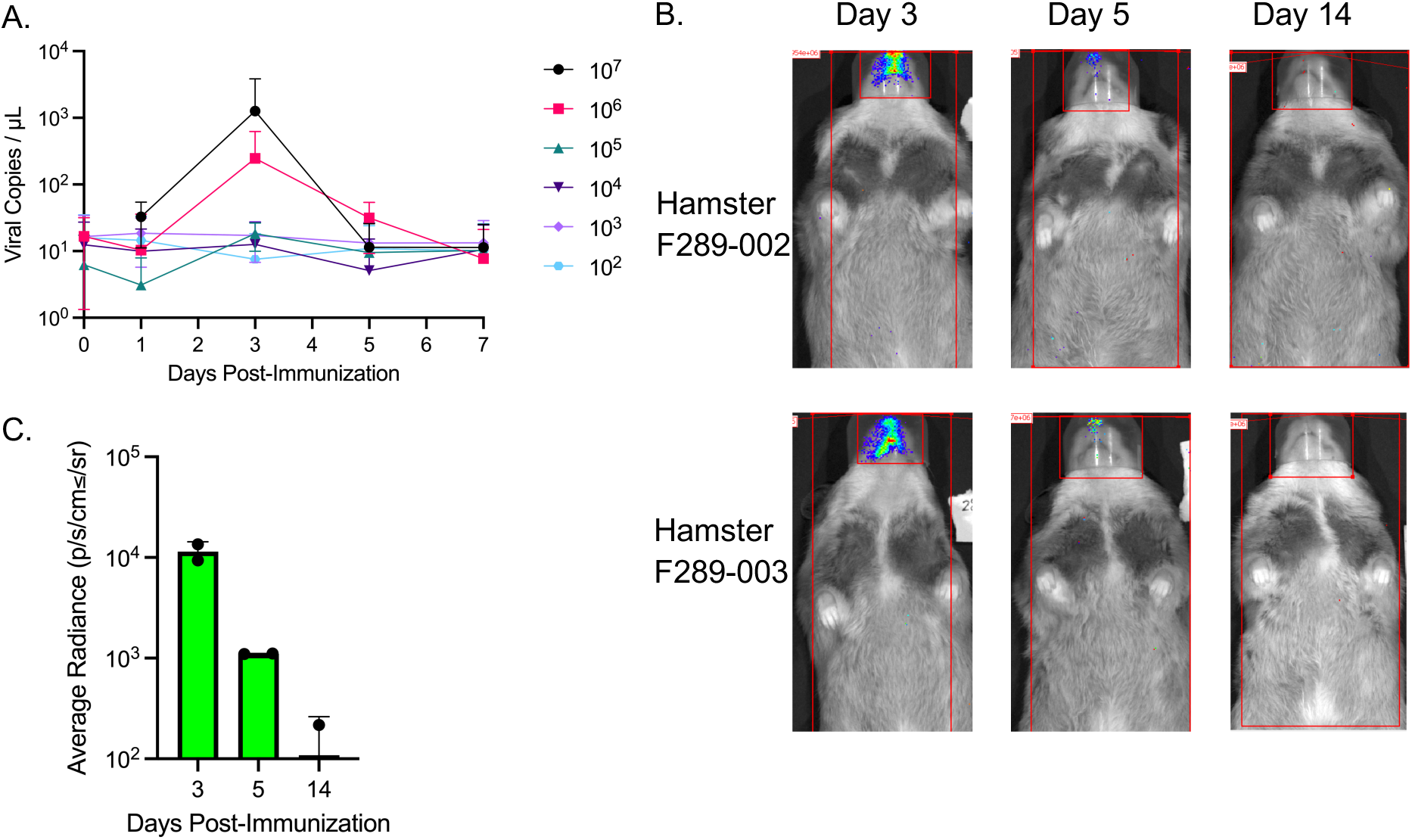
Ad4 replication and transgene expression over time after IN inoculation. (A) Viral copies/µL recovered from hamster oropharyngeal swabs was measured by qPCR following inoculation with 10^2^-10^7^ IFU of Ad4 on Day 0. (B) Bioluminescence measured in anesthetized male hamsters after IN inoculation with 6.8x10^6^ IFU of Ad4 expressing luciferase on Day 0. Average radiance was calculated after subtracting the average radiance of the nose of an uninoculated hamster.

**Supplementary Figure 1.**
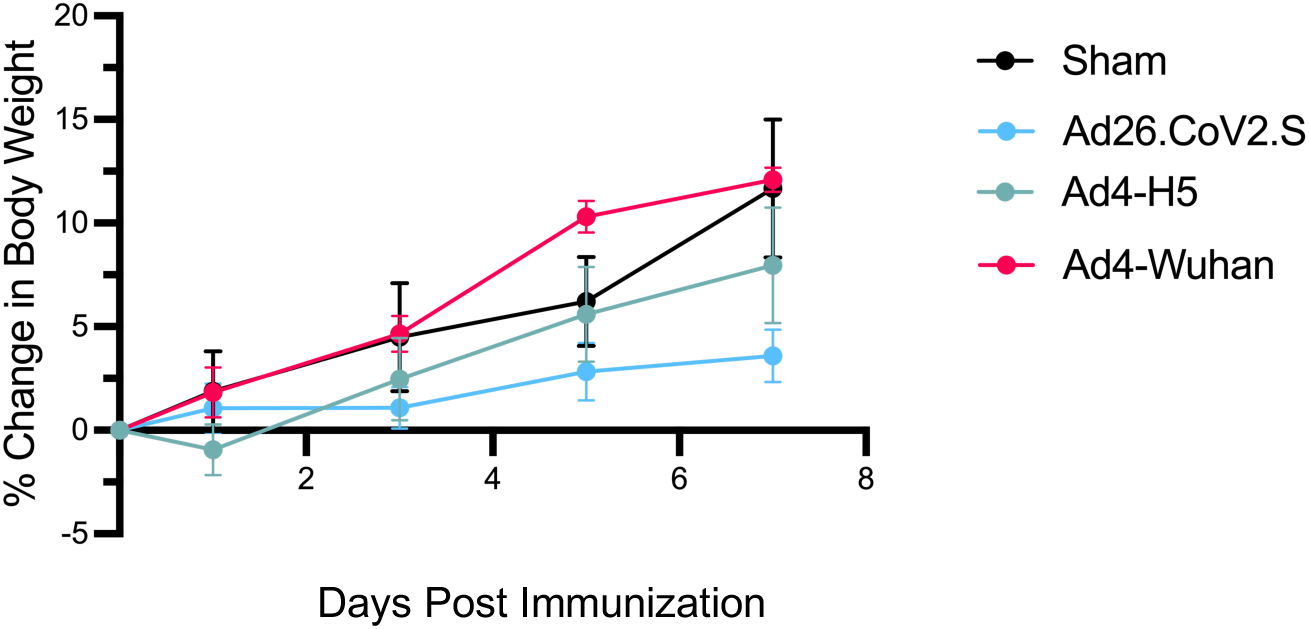
Immunization with Ad4-constructs alone does not significantly impact hamster weight. Longitudinal weight change of hamsters administered IM sham, IM Ad26.CoV2.S, IN Ad4-H5, or Ad4-Wuhan.

We then made Ad4 recombinants expressing Spike antigens from variants of concern (Wuhan (B), Beta (B.1.351), Delta (B.1.617.2), Gamma (P.1)), a consensus of Wuhan and Beta, and another expressing influenza virus hemagglutinin H5 (Ad4-H5) as a negative control, and intranasally administered 10^7^ IFU or a saline (sham) control. The Ad26 based vaccine (Ad26.CoV2.S; Johnson and Johnson) was administered IM as a comparator. To examine the systemic and mucosal humoral immune responses to SARS-CoV-2 following immunization, Syrian hamsters were vaccinated with 10^7^ IFU of the Ad4 constructs or controls and binding antibodies (IgG, IgA, and IgM) in hamster nasal wash, bronchoalveolar lavage, and serum samples were measured against an array of SARS-CoV-2 antigens at 56 days following the first immunization. As expected, no binding to SARS-CoV-2 nucleocapsid was detected in samples from nasal wash, BAL, or serum. There was only low-level cross-reactive binding to the Omicron receptor binding domain (RBD) or SARS-CoV-1 S. Hamsters immunized with either IM Ad26.CoV2.S or each IN Ad4-Spike recombinants had higher SARS-CoV-2-specific IgG, IgA, and IgM antibodies in the BAL relative to those given the sham vaccination or Ad4-H5. However, only the IN Ad4-Spike recombinants induced such binding antibody responses in the nasal wash (Figure 2, p<0.01, all comparisons). These data suggested that unlike the IM administered Ad26.CoV2.S, IN administration of the Ad4-Wuhan construct induced both systemic and mucosal binding antibodies.

**Figure 2.**
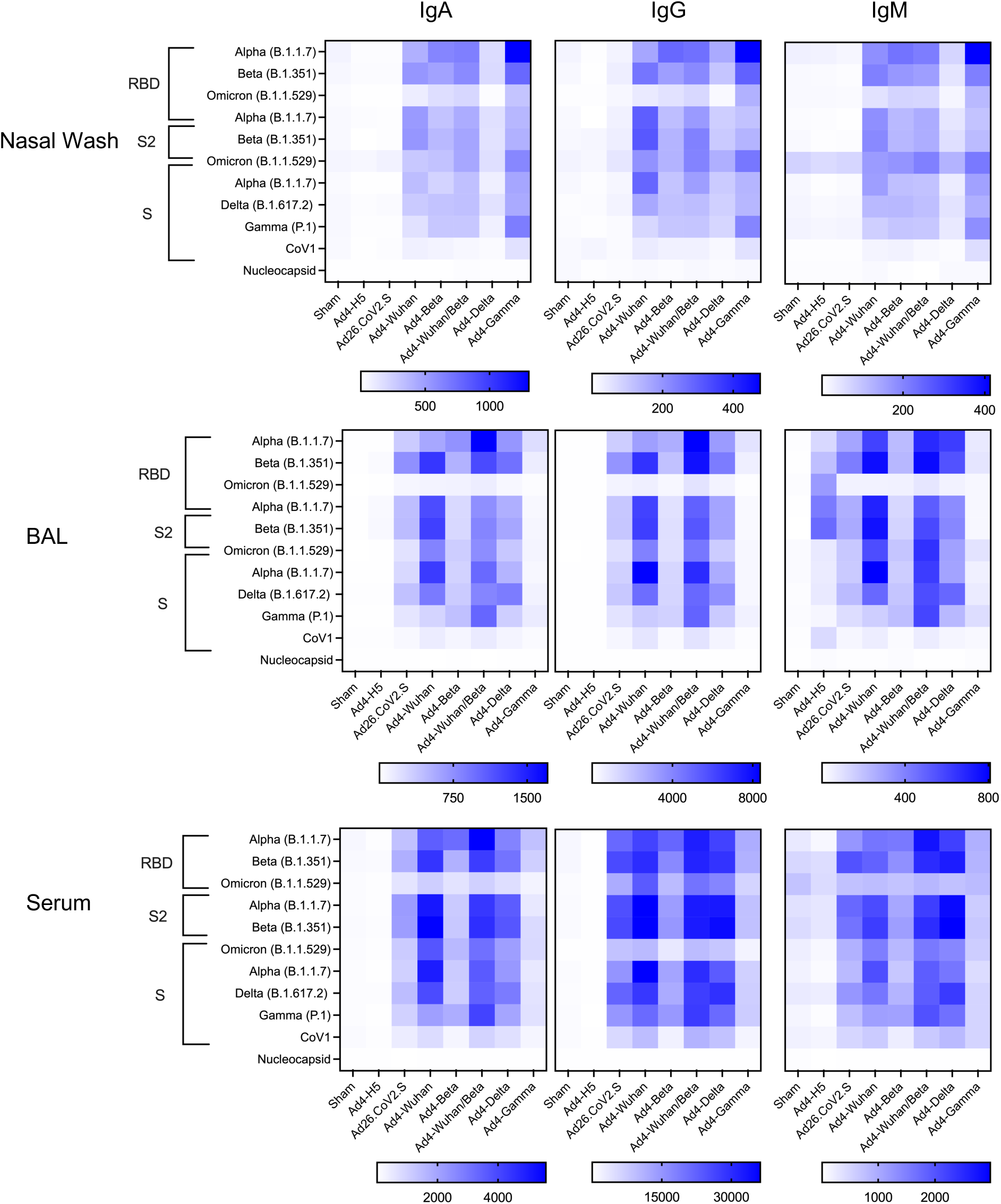
IN Ad4-SARS-CoV-2 induces binding antibodies on the nasal mucosa. Heat map depicting binding IgA (left), IgG (center) and IgM (right) responses observed in hamster nasal wash (top), BAL (center), and serum (bottom) for each immunization group (columns) against the indicated SARS-CoV antigens (rows), including Spike (S), the S2 and receptor binding domain (RBD), and Nucleocapsid protein. Mean response values in each group of 6 animals is graphed.

To further assess the immunogenicity and efficacy of immune responses induced by IN immunization with the Ad4-Spike recombinant vaccines, we then performed a challenge study that included IN Ad4-Wuhan (B), -Delta (B.1.617.2), and -Omicron (BA.2.75.2). We also included Ad26CoV2.S and mRNA-1273 (Moderna) immunized animals as comparators, and Ad4-H5 vaccinated hamsters as a negative control. Animals were bled and challenged at an early (day 56) or late (day 174) timepoint (Supplementary Fig. 2). The later timepoint permitted an assessment of durability of the response and an evaluation of efficacy at a timepoint when the innate immune response had subsided and the CD8^+^ T cells have contracted into a memory response. Prior to challenge, the serum neutralizing antibody titers were measured against a panel of pseudotyped lentiviruses expressing spike from Wuhan (B.1.1), Delta (B.1.617.2), or Omicron (B.1.1.529 and BA.2.75.2) lineages (Fig. 3). When tested against the Wuhan pseudovirus, immunization with IN Ad4-Wuhan or Ad4-Delta constructs induced neutralizing antibodies at day 56 comparable to IM mRNA-1273 (Median ID_50_: Ad4-Wuhan = 1684.5, Ad4-Delta = 992.9, mRNA-1273 = 413.5). However, the titers induced by Ad4-Wuhan or Ad4-Delta were significantly higher than Ad26.CoV2.S (Ad26.CoV2.S = 210.9; p=0.007 and 0.033, respectively). A similar pattern was observed in neutralization of the Delta pseudovirus (Median ID_50_: Ad4-Wuhan = 629.2, Ad4-Delta = 451.8, mRNA-1273 = 306, Ad26.CoV2.S = 211.1; Ad4-Wuhan vs Ad26.CoV2.S p=0.002; Ad4-Delta vs Ad26.CoV2.S p=0.02). As expected, the serum neutralization against Omicron variants was more modest. Animals immunized with the IN Ad4-Delta, but not Ad4-Wuhan, had significantly higher neutralization titers against the Omicron B.1.529 pseudovirus compared to IM Ad26.CoV2.S or mRNA-1273 vaccines (Median ID_50_: Ad4-Delta Median ID_50_ = 276, Ad4-Wuhan = 108.3, mRNA-1273 = 29, Ad26.CoV2.S = 43.2; p=0.030, both comparisons) suggesting greater breadth. The serum neutralization potency induced by Ad4-Wuhan or Ad4-Delta against the Omicron BA.2.75.2 pseudovirus was low and comparable to those induced by the IM Ad26.CoV2.S or mRNA-1273 vaccines (Median ID_50_: Ad4-Delta = 50.4, Ad4-Wuhan = 40, mRNA-1273 = 29, Ad26.CoV2.S = 37.9; p=0.001, both comparisons). This lack of heterologous neutralization may be partially attributed to the relatively large antigenic distance between the Wuhan or Delta lineages and those of Omicron BA.2.75.2^15,34^. However, the Ad4-BA.2.75.2 construct did induce higher neutralizing antibody titers when tested against the Omicron B.1.1.529 pseudovirus compared to mRNA-1273 or Ad26.CoV2.S (Median ID_50_: Ad4-BA.2.75.2 = 1940, mRNA-1273 = 29, Ad26.CoV2.S = 43; p<0.001 and p=0.001, respectively), despite failing to induce detectable neutralization against the Wuhan or Delta pseudoviruses (Median ID_50_: Ad4-BA.2.75.2 = 29 and 29, respectively) (Fig. 3). A similar pattern was observed against the Omicron BA.2.75.2 pseudovirus (Median ID50: Ad4-BA.2.75.2 = 1247, mRNA-1273 = 29, Ad26.CoV2.S = 38; p<0.001 for comparisons with Ad26.CoV.2 or mRNA-1273).

**Figure 3.**
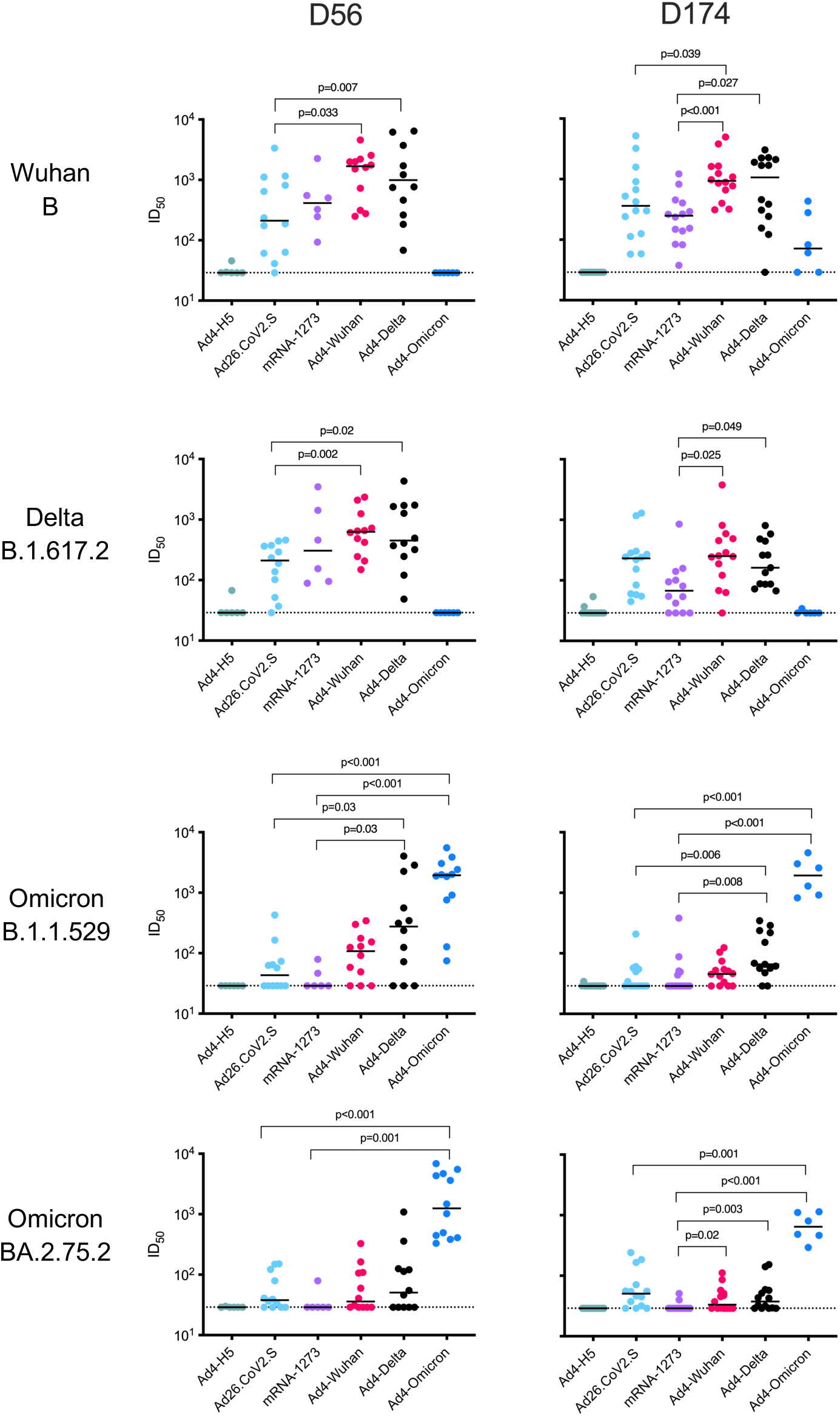
IN immunization with Ad4-Spike induces a systemic neutralizing antibodies. ID50 values for each hamster against the indicated pseudovirus at 56 (left) or 174 (right) days after immunization are indicated with dots. Median ID_50_ values are indicated with a black line. The dotted line indicates the limit of detection of 30.

**Supplementary Figure 2.**
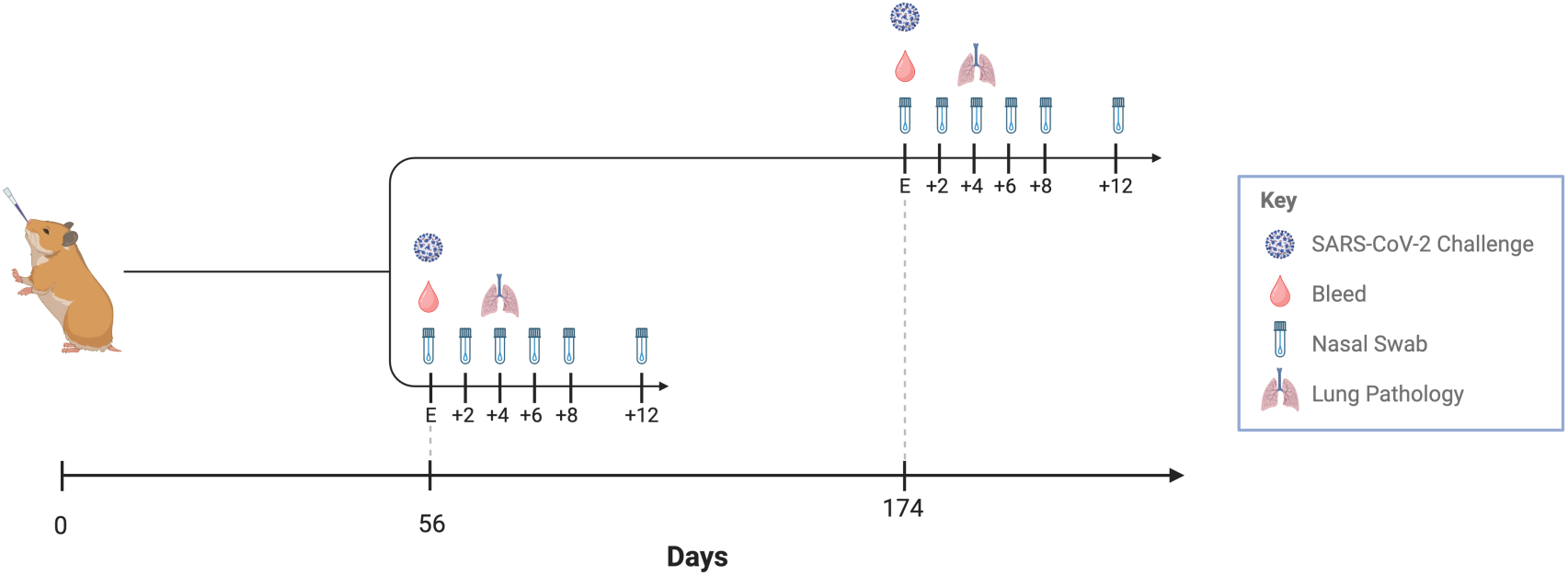
Challenge study outline. Hamsters received IN or IM vaccines on day 0 and then challenged on day 56 or day 174 with either SARS-CoV-2 Delta or Omicron. Samples were then taken at the indicated timepoints.

Rapid waning of serum neutralizing antibody titers post-vaccination is an undesirable feature of mRNA vaccines^35-37^. This contrasts with our experience with replicating Ad4 expressing hemagglutinin H5 in humans where serum neutralizing antibodies were unchanged for up to 5 years after IN vaccination^25,26^. To characterize the durability of the immune responses induced by the Ad4-Spike immunogens, we further assessed serum neutralization at 174 days (Fig. 3). Neutralizing antibody titers against the Wuhan pseudovirus remained higher in Ad4-Wuhan immunized animals compared to Ad26.CoV2.S (Median ID_50_: Ad4-Wuhan = 936, Ad26.CoV2.S = 362; p=0.039) and both Ad4-Wuhan and Ad4-Delta groups were higher than those immunized with mRNA-1273 (Median ID_50_: Ad4-Delta = 1071; mRNA-1273 = 250; p<0.001 and p= 0.027, respectively). Likewise, serum neutralizing antibody titers for Ad4-Wuhan and Ad4-Delta immunized groups against Delta pseudovirus were comparable with Ad26.CoV2.S group but were significantly higher compared to mRNA-1273 group (Median ID_50_: Ad4-Wuhan = 251.4, Ad4-Delta = 162, Ad26.CoV2.S = 231; mRNA-1273 = 68; p=0.025 and 0.049, respectively). Neutralizing antibody titers against the Omicron B.1.1.529 and BA.2.75.2 pseudoviruses remained low at 174 days. In contrast, neutralizing antibody titers in animals immunized with the IM Ad26.CoV2.S vaccine or IN Ad4-Spike constructs did not change by 174 DPI when tested against the Wuhan pseudovirus (Median ID_50_: Ad26.CoV2.S = 362; Ad4-Wuhan ID_50_ = 936; Ad4-Delta ID_50_ = 1071). A similar pattern was observed against the Delta pseudovirus (Median ID_50_: Ad26.CoV2.S = 231; Ad4-Wuhan ID_50_ = 251; Ad4-Delta ID_50_ = 162). The neutralizing antibody responses in animals that received the Ad4-BA.2.75.2 vaccine were similarly durable against the Omicron B.1.1.529 pseudovirus (Median ID_50_: Ad4-BA.2.75.2 = 1940 day 56, 1934 day 174) and BA.2.75.2 (Median ID_50_: Ad4-BA.2.75.2 = 1247 day 56, 642 day 174).

Systemic and mucosal binding antibodies were also measured at day 180 in a subset of animals immunized with mRNA-1273, Ad4-Delta, or Ad4-Omicron BA.2.75.2. Each of these vaccines induced serum binding antibodies, dominated by those directed to the relatively conserved S2 subunit of the Spike protein (Supplementary Fig. 3). On the whole IgA, IgG, and IgM antibodies that bound the receptor binding domain (RBD) or S were higher in IN Ad4-Delta or Ad4-Omicron-immunized animals compared to mRNA-1273-immunized animals (Supplementary Table 1). At this late timepoint the nasal IgA, IgG, and IgM antibodies were very low among the vaccine groups consistent with the waning of nasal responses that has been observed in humans^9,38,39^.

**Supplementary Figure 3.**
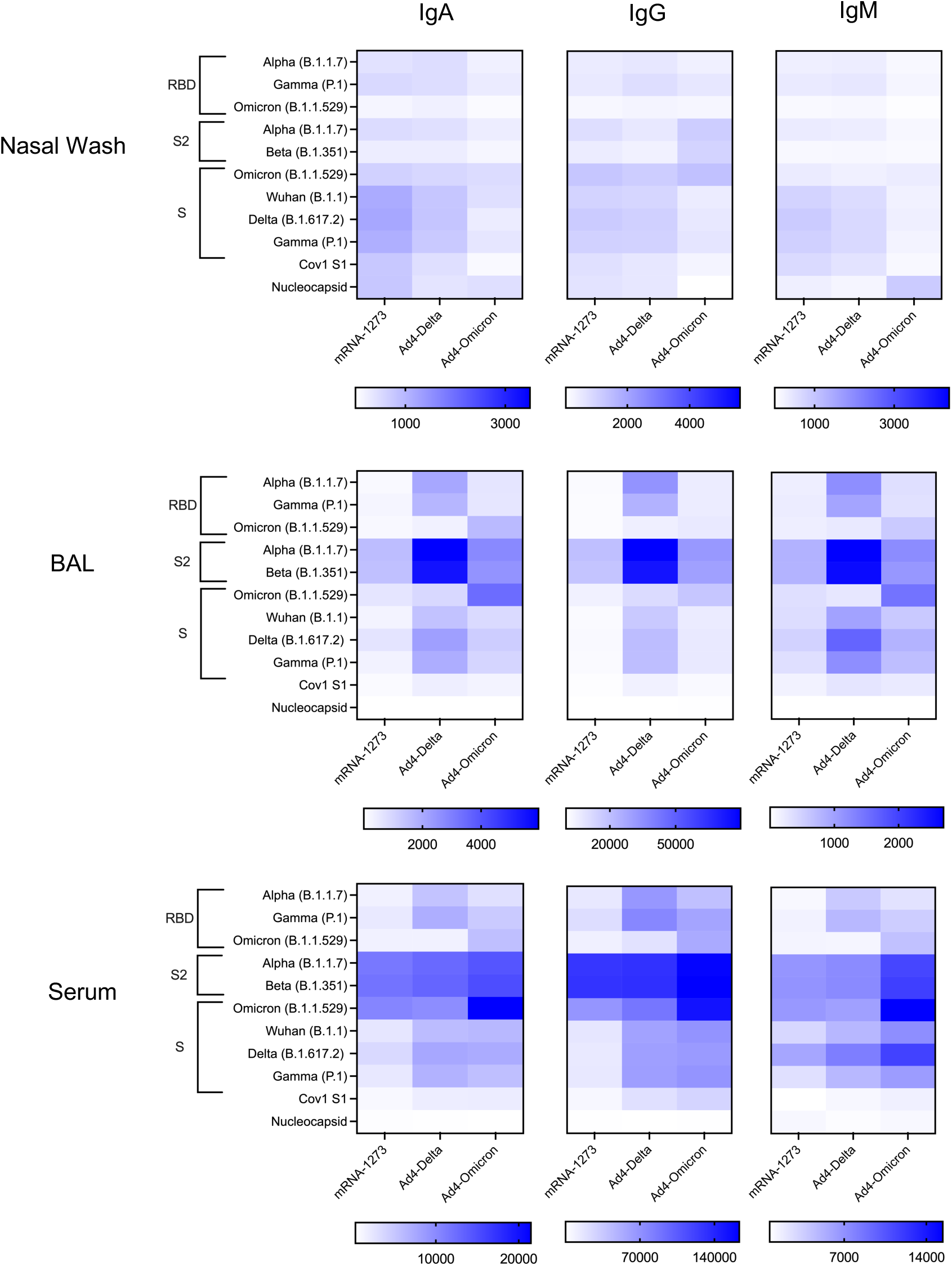
IN Ad4-Spike binding antibody responses from the D180 nasal mucosa. Binding Ig characterization in hamster nasal wash, BAL, and serum against the indicated SARS-CoV antigens, including Spike (S) and Spike subunits, the receptor binding domain (RBD), and Nucleocapsid proteins.

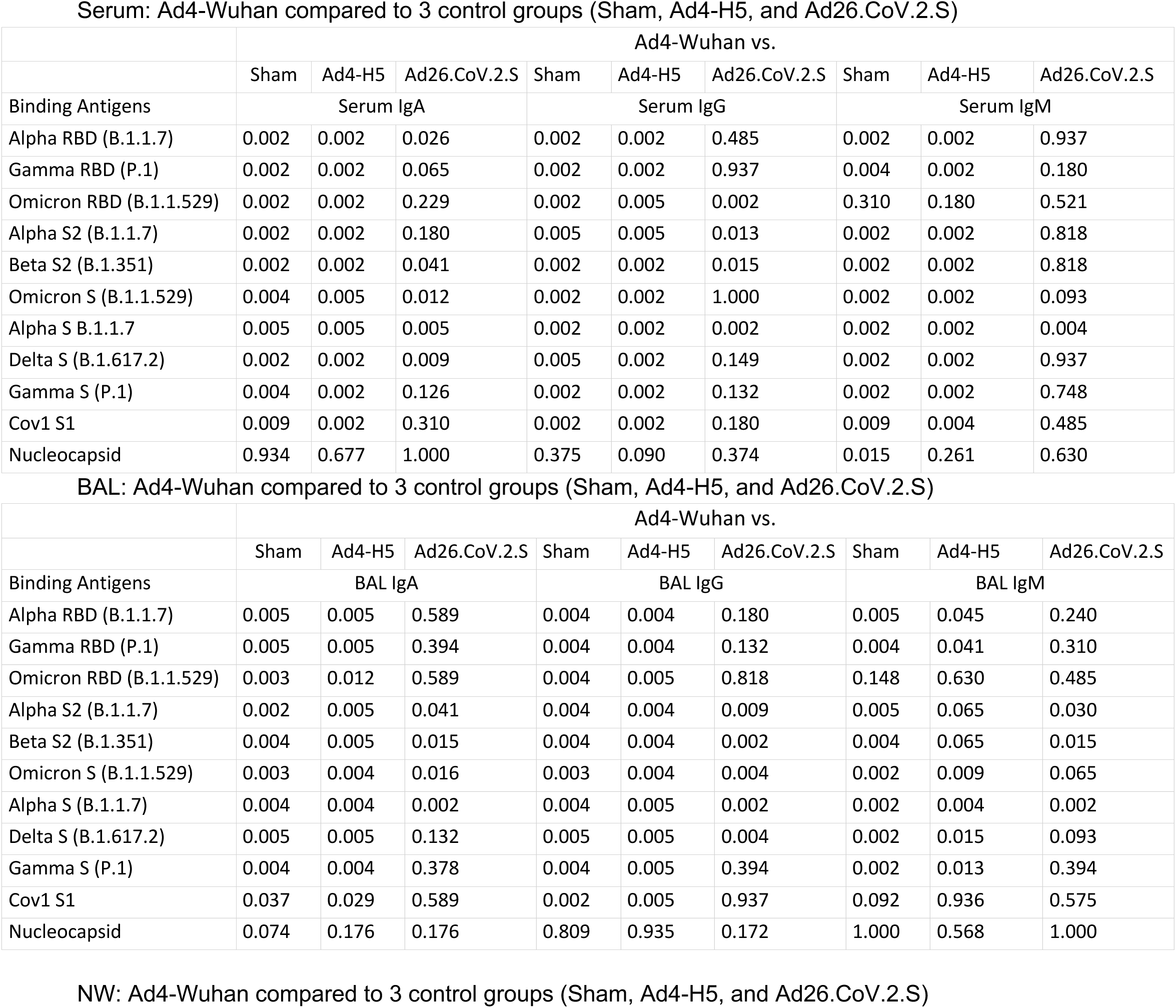
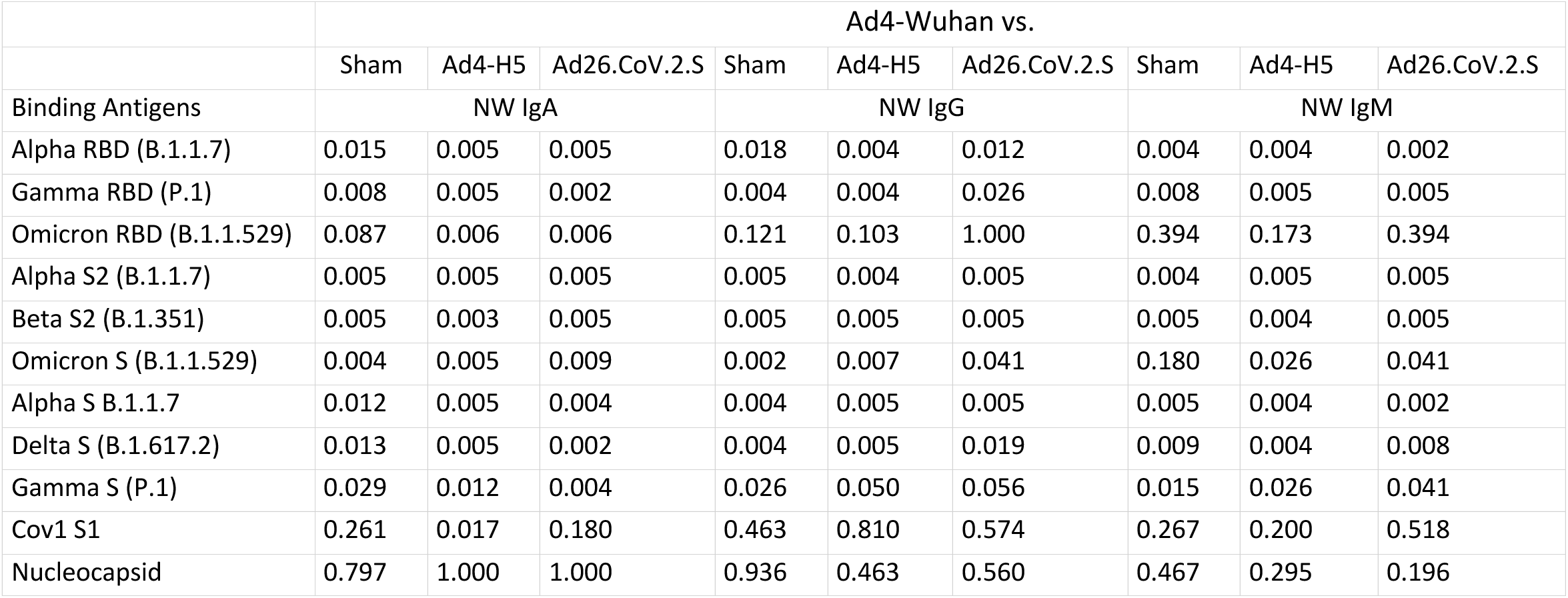

|  | Ad4-Wuhan vs. |  |  |  |  |  |  |  |  |
| --- | --- | --- | --- | --- | --- | --- | --- | --- | --- |
|  | Sham | Ad4-H5 | Ad26.CoV.2.S | Sham | Ad4-H5 | Ad26.CoV.2.S | Sham | Ad4-H5 | Ad26.CoV.2.S |
| Binding Antigens | Serum IgA |  |  | Serum IgG |  |  | Serum IgM |  |  |
| Alpha RBD (B.1.1.7) | 0.002 | 0.002 | 0.026 | 0.002 | 0.002 | 0.485 | 0.002 | 0.002 | 0.937 |
| Gamma RBD (P.1) | 0.002 | 0.002 | 0.065 | 0.002 | 0.002 | 0.937 | 0.004 | 0.002 | 0.180 |
| Omicron RBD (B.1.1.529) | 0.002 | 0.002 | 0.229 | 0.002 | 0.005 | 0.002 | 0.310 | 0.180 | 0.521 |
| Alpha S2 (B.1.1.7) | 0.002 | 0.002 | 0.180 | 0.005 | 0.005 | 0.013 | 0.002 | 0.002 | 0.818 |
| Beta S2 (B.1.351) | 0.002 | 0.002 | 0.041 | 0.002 | 0.002 | 0.015 | 0.002 | 0.002 | 0.818 |
| Omicron S (B.1.1.529) | 0.004 | 0.005 | 0.012 | 0.002 | 0.002 | 1.000 | 0.002 | 0.002 | 0.093 |
| Alpha S B.1.1.7 | 0.005 | 0.005 | 0.005 | 0.002 | 0.002 | 0.002 | 0.002 | 0.002 | 0.004 |
| Delta S (B.1.617.2) | 0.002 | 0.002 | 0.009 | 0.005 | 0.002 | 0.149 | 0.002 | 0.002 | 0.937 |
| Gamma S (P.1) | 0.004 | 0.002 | 0.126 | 0.002 | 0.002 | 0.132 | 0.002 | 0.002 | 0.748 |
| Cov1 S1 | 0.009 | 0.002 | 0.310 | 0.002 | 0.002 | 0.180 | 0.009 | 0.004 | 0.485 |
| Nucleocapsid | 0.934 | 0.677 | 1.000 | 0.375 | 0.090 | 0.374 | 0.015 | 0.261 | 0.630 |

|  | Ad4-Wuhan vs. |  |  |  |  |  |  |  |  |
| --- | --- | --- | --- | --- | --- | --- | --- | --- | --- |
|  | Sham | Ad4-H5 | Ad26.CoV.2.S | Sham | Ad4-H5 | Ad26.CoV.2.S | Sham | Ad4-H5 | Ad26.CoV.2.S |
| Binding Antigens | BAL IgA |  |  | BAL IgG |  |  | BAL IgM |  |  |
| Alpha RBD (B.1.1.7) | 0.005 | 0.005 | 0.589 | 0.004 | 0.004 | 0.180 | 0.005 | 0.045 | 0.240 |
| Gamma RBD (P.1) | 0.005 | 0.005 | 0.394 | 0.004 | 0.004 | 0.132 | 0.004 | 0.041 | 0.310 |
| Omicron RBD (B.1.1.529) | 0.003 | 0.012 | 0.589 | 0.004 | 0.005 | 0.818 | 0.148 | 0.630 | 0.485 |
| Alpha S2 (B.1.1.7) | 0.002 | 0.005 | 0.041 | 0.004 | 0.004 | 0.009 | 0.005 | 0.065 | 0.030 |
| Beta S2 (B.1.351) | 0.004 | 0.005 | 0.015 | 0.004 | 0.004 | 0.002 | 0.004 | 0.065 | 0.015 |
| Omicron S (B.1.1.529) | 0.003 | 0.004 | 0.016 | 0.003 | 0.004 | 0.004 | 0.002 | 0.009 | 0.065 |
| Alpha S (B.1.1.7) | 0.004 | 0.004 | 0.002 | 0.004 | 0.005 | 0.002 | 0.002 | 0.004 | 0.002 |
| Delta S (B.1.617.2) | 0.005 | 0.005 | 0.132 | 0.005 | 0.005 | 0.004 | 0.002 | 0.015 | 0.093 |
| Gamma S (P.1) | 0.004 | 0.004 | 0.378 | 0.004 | 0.005 | 0.394 | 0.002 | 0.013 | 0.394 |
| Cov1 S1 | 0.037 | 0.029 | 0.589 | 0.002 | 0.005 | 0.937 | 0.092 | 0.936 | 0.575 |
| Nucleocapsid | 0.074 | 0.176 | 0.176 | 0.809 | 0.935 | 0.172 | 1.000 | 0.568 | 1.000 |
|  | Sham | Ad4-H5 | Ad26.CoV.2.S | Sham | Ad4-H5 | Ad26.CoV.2.S | Sham | Ad4-H5 | Ad26.CoV.2.S |
| Binding Antigens | NW IgA |  |  | NW IgG |  |  | NW IgM |  |  |
| Alpha RBD (B.1.1.7) | 0.015 | 0.005 | 0.005 | 0.018 | 0.004 | 0.012 | 0.004 | 0.004 | 0.002 |
| Gamma RBD (P.1) | 0.008 | 0.005 | 0.002 | 0.004 | 0.004 | 0.026 | 0.008 | 0.005 | 0.005 |
| Omicron RBD (B.1.1.529) | 0.087 | 0.006 | 0.006 | 0.121 | 0.103 | 1.000 | 0.394 | 0.173 | 0.394 |
| Alpha S2 (B.1.1.7) | 0.005 | 0.005 | 0.005 | 0.005 | 0.004 | 0.005 | 0.004 | 0.005 | 0.005 |
| Beta S2 (B.1.351) | 0.005 | 0.003 | 0.005 | 0.005 | 0.005 | 0.005 | 0.005 | 0.004 | 0.005 |
| Omicron S (B.1.1.529) | 0.004 | 0.005 | 0.009 | 0.002 | 0.007 | 0.041 | 0.180 | 0.026 | 0.041 |
| Alpha S B.1.1.7 | 0.012 | 0.005 | 0.004 | 0.004 | 0.005 | 0.005 | 0.005 | 0.004 | 0.002 |
| Delta S (B.1.617.2) | 0.013 | 0.005 | 0.002 | 0.004 | 0.005 | 0.019 | 0.009 | 0.004 | 0.008 |
| Gamma S (P.1) | 0.029 | 0.012 | 0.004 | 0.026 | 0.050 | 0.056 | 0.015 | 0.026 | 0.041 |
| Cov1 S1 | 0.261 | 0.017 | 0.180 | 0.463 | 0.810 | 0.574 | 0.267 | 0.200 | 0.518 |
| Nucleocapsid | 0.797 | 1.000 | 1.000 | 0.936 | 0.463 | 0.560 | 0.467 | 0.295 | 0.196 |

Hamsters were then challenged intranasally with SARS-CoV-2 Delta (B.1.617.2) or Omicron (B.1.1.529) at 56 or 174 days to further characterize the protective efficacy of the Ad4-Spike constructs against SARS-CoV-2 infection. When challenged with SARS-CoV-2 Delta at day 56, hamsters immunized with each of the SARS-CoV-2 vaccines had less weight loss compared to those that received Ad4-H5 (p<0.001 for all comparisons) (Fig. 4). However, IN Ad4-Wuhan immunized animals had less weight loss compared to mRNA-1273 or Ad26.CoV2.S (p=0.043 and 0.009, respectively). A similar trend was observed for Ad4-Delta (p=0.011 and 0.003, respectively). These weight loss trends between groups were similar to those observed when hamsters were challenged with SARS-CoV-2 Delta at day 174. IN Ad4-Wuhan-immunized animals had less weight loss compared to mRNA-1273 or Ad26.CoV2.S (p=0.002 and <0.001, respectively). And a similar trend was observed for Ad4-Delta compared to mRNA-1273 or Ad26.CoV2.S (p=0.004 and p=0.002, respectively). Differences in weight loss between groups that were challenged with SARS-CoV-2 Omicron on day 56 did not achieve significance, likely because this variant is not as virulent as Delta in hamsters (Fig. 4B)^34^. Following SARS-CoV-2 Omicron challenge at day 174, Ad4-Wuhan and Ad4-Delta-immunized animals had significantly less weight loss compared to the Ad4-H5 control group (p= 0.002 and 0.005, respectively) as did Ad26.CoV2.S (p=0.043, both comparistons) although these differences were modest.

**Figure 4.**
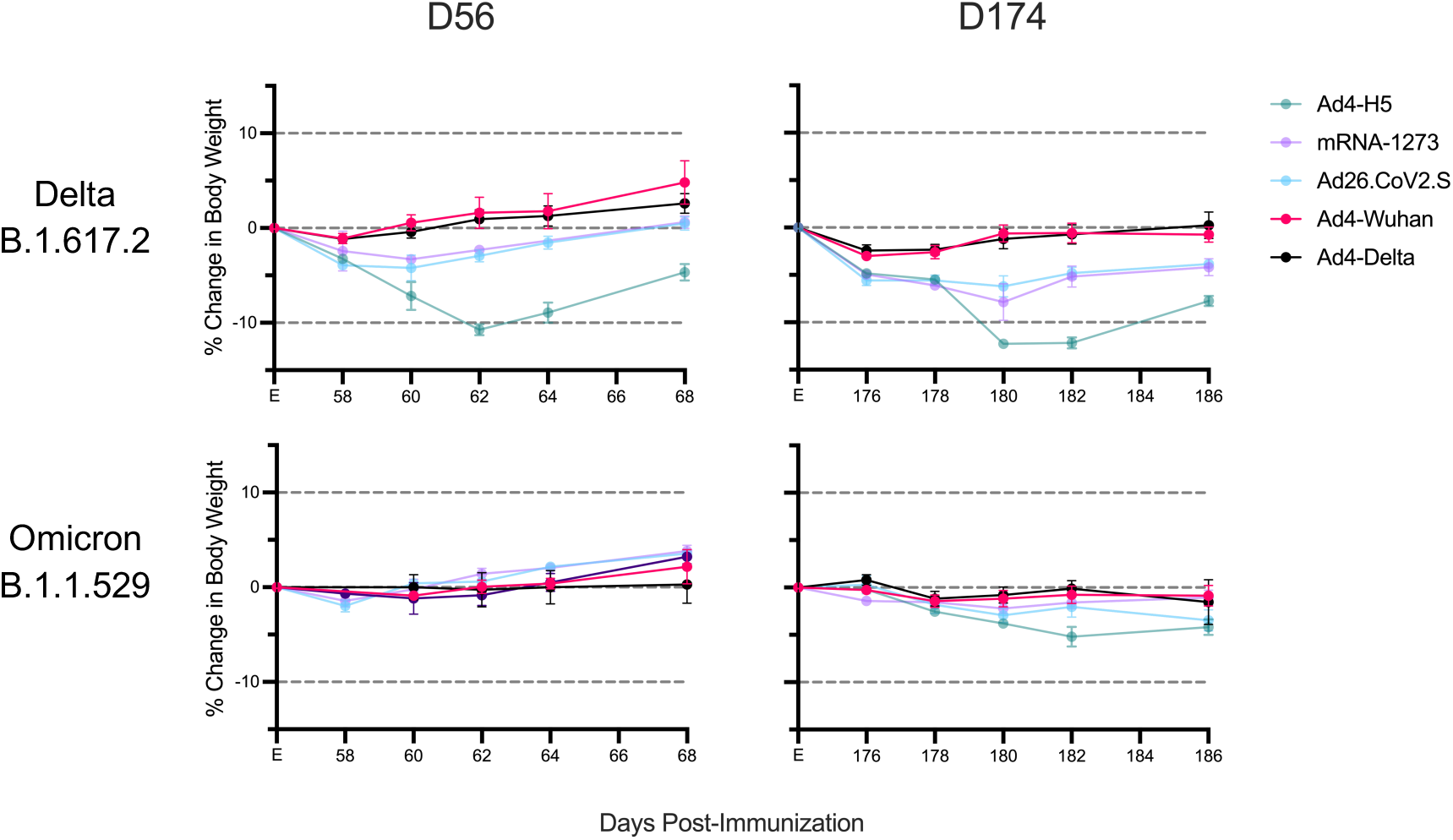
IN Ad4-Spike immunization limits weight loss following SARS-CoV-2 challenge. Average percent body weight relative to the day of challenge, at either 56 days or 174 days following the first immunization. Dots represent median and bars represent standard error.

To investigate the impact of IN immunization on viral replication, subgenomic RNA (sgRNA) was extracted from oropharyngeal swabs, polymerase chain reaction amplified, and measured up to 14 days post-challenge. When hamsters were challenged with SARS-CoV-2 Delta at 56 days, each of the SARS-CoV-2 vaccines reduced viral replication compared to Ad4-H5 (Median AUC (copies/ml): Ad26.CoV2.S = 6.9 x 10^7^; mRNA1273 = 2.1 x 10^7^; Ad4-Wuhan = 3.4 x 10^7^; Ad4-Delta = 3.6 x 10^7^; Ad4-H5 = 2.5 x 10^8^ p<0.011p<0.001, p=0.003, and p=0.002, respectively), but were similar in protective efficacy (Fig. 5). Ad4-Wuhan-immunized animals had significant reductions in Omicron challenge virus replication compared to Ad4-H5-, Ad26.CoV2.S-or mRNA-1273-immunized animals (Median AUC: Ad4-Wuhan = 1.4 x 10^6^; Ad4-H5 = 1.1 x 10^7^; Ad26.CoV2.S = 1.6 x 10^7^; mRNA-1273 = 7.9 x 10^7^; p=0.002, p=0.001, and p=0.023, respectively), consistent with greater breadth of the response. Ad4-Delta immunized animals also had lower viral replication compared to those immunized with Ad26.CoV2.S or Ad4-H5 (Median AUC: Ad4-Delta = 2.1 x 10^6^, Ad26.CoV2.S = 1.6 x 10^7^, Ad4-H5 = 1.1 x 10^7^; p=0.002 p=0.011, respectively). After SARS-CoV-2 Delta challenge at day 174, animals immunized with Ad4-Wuhan had lower levels of SARS-CoV-2 Delta replication compared to those immunized with Ad4-H5 or mRNA-1273 (Median AUC: Ad4-Wuhan = 5.9 x 10^6^; Ad4-H5 = 3.5 x 10^7^; mRNA-1273 = 6.2 x 10^7^; p<0.001 for both comparisons). Ad4-Delta immunized animals also had lower levels of viral replication compared to both Ad26.CoV2.S or mRNA-1273 (Median AUC: Ad4-Delta = 4.2 x 10^6^; Ad26.CoV2.S = 1 x 10^7^; p=0.043 and <0.001, respectively). The trend for greater breadth of protection induced by the Ad4-S recombinants against the SARS-CoV-2 Omicron virus was maintained at day 174. Whereas neither Ad26.CoV2.S nor mRNA-1273 significantly reduced SARS-CoV-2 Omicron replication relative to Ad4-H5 (p>0.5, both comparisons), Ad4-Wuhan reduced replication relative to both Ad26.CoV2.S and mRNA-1273 (Median AUC: Ad4-Wuhan = 4.9 x 10^6^; Ad4-H5 = 4.0 x 10^7^; Ad26.CoV2.S = 4.5 x 10^7^; mRNA-1273 = 3.0 x 10^7^; p<0.001 and p=0.004, respectively). Similarly, Ad4-Delta reduced SARS-CoV-2 Omicron replication relative to these two IM vaccines (Median AUC: Ad4-Delta = 1.5 x 10^6^; p<0.001, both comparisons). Taken together these results suggest that both Ad4-Wuhan and Ad4-Delta induced greater breadth and durability of protection relative to Ad26.CoV2.S and mRNA-1273.

**Figure 5.**
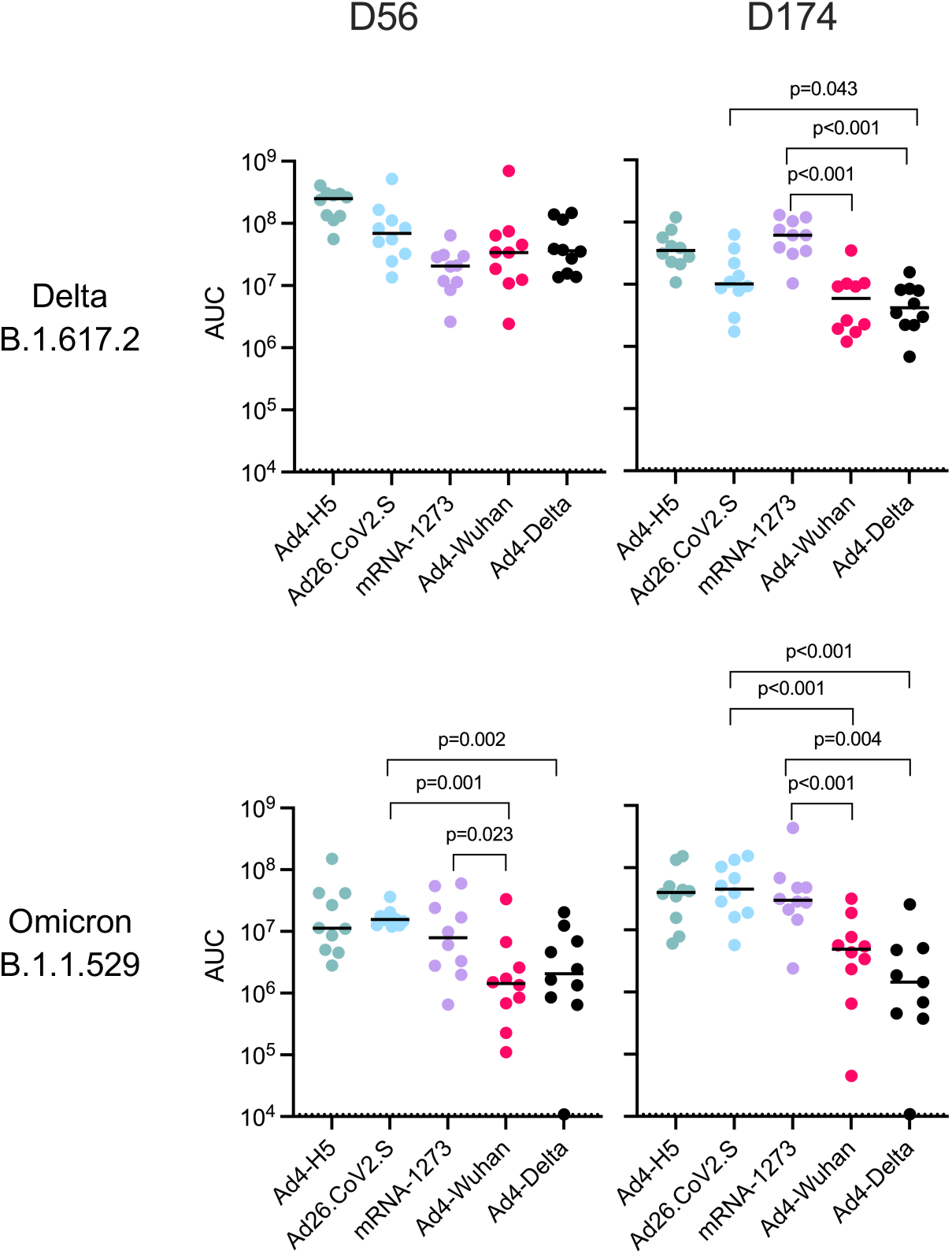
IN immunization with Ad4-Spike reduces challenge virus replication in the upper respiratory tract. Subgenomic RNA was detected in hamster oropharyngeal swabs following challenge at day 56 or day 174. For each hamster, the number of viral copies in each swab was plotted over time, and the area under the curve (AUC) that resulted from these plots was calculated. The dotted line indicates the AUC limit of detection over 12 days (10,800 copies). The median AUC value for each group is represented as a black line.

To evaluate the impact of immunization on lower respiratory tract disease, four hamsters per group were sacrificed at four days after SARS-CoV-2 challenge at days 56 or 174 to evaluate pulmonary pathology. Pathology slides were graded in a blinded manner for SARS-CoV-2 antigen by immunohistochemistry (IHC), inflammation severity, and percent of lung tissue affected. At day 60 (four days after challenge), the animals immunized with IN Ad4-Wuhan or Ad4-Delta had lower pathology scores than those that received Ad4-H5 (IHC alveoli, bronchial epithelium, percent lung affected, and severity; p<0.05 for all comparisons)(Fig. 6). Most scores of animals immunized with IN Ad4-S recombinants were below those of IM Ad26.CoV2.S (IHC alveoli, IHC bronchial epithelium, percent lung affected;p<0.05 for each comparison), but similar to those of IM mRNA-1273 (p>0.05 for all comparisons). Following challenge with SARS-CoV-2 Omicron (B.1.1.529) at day 60 pathology scores were lower overall for IN Ad4-S recombinant immunized groups compared to IM Ad26.CoV2.S or mRNA-1273,, consistent with the lack of weight loss observed in this study and in prior work ^40^. Although the trends for a reduction in pathology in Ad4-Wuhan immunized animals after SARS-CoV-2 Omicron challenge were similar to those of SARS-CoV-2 Delta challenged animals, only the IHC scores in the alveoli were significantly different than Ad4-H5 (p=0.018) and Ad26.CoV2.S (p=0.020). Following SARS-CoV-2 Delta challenge at day 174, Ad4-Wuhan immunized animals had scores that were below those that received Ad4-H5 (IHC alveoli p=0.027, IHC bronchial p=0.025, pneumonia severity p=0.040), and similar to those that received Ad26.CoV2.S (p>0.05 for all comparisons), but below those that received mRNA-1273 (IHC alveoli p=0.025, IHC bronchial 0.023, pneumonia severity p=0.026). Similarly, following SARS-CoV-2 Omicron challenge at day 174, Ad4-Wuhan immunized animals had lower pathology scores compared to those immunized with mRNA-1273 (IHC bronchial p=0.020; percent lung affected p= 0.026). Overall, the Ad4-Wuhan vaccine offered the most robust protection from pulmonary pathology following SARS-CoV-2 challenge, with the lowest average scores at both time points for both challenge viruses.

**Figure 6.**
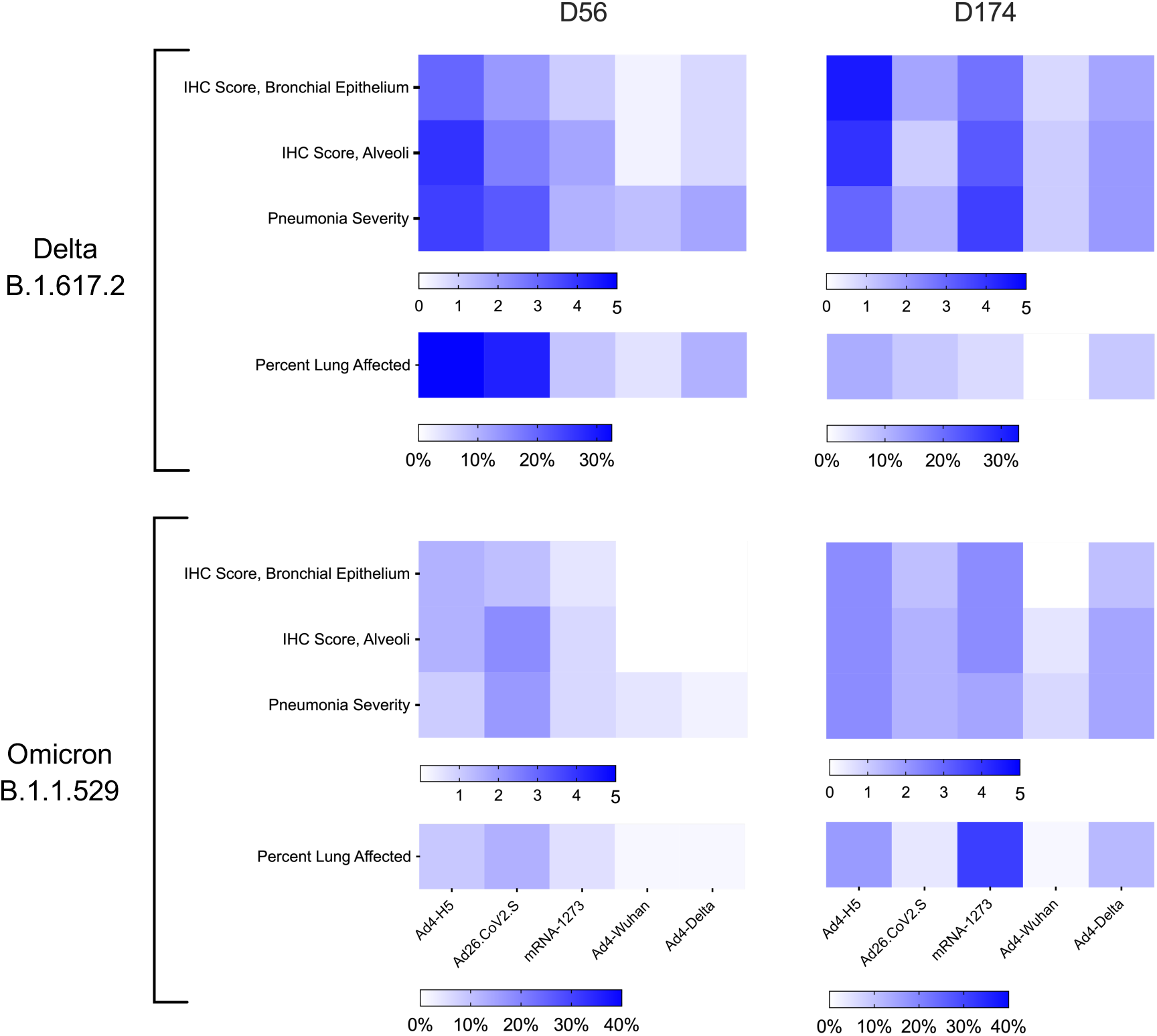
IN Immunization with Ad4-Wuhan reduces lung pathology following SARS-CoV-2 challenge. Four hamsters from each group were sacrificed four days after challenge (day 60 or 178) to assess disease severity in the lungs. Scores ranging from 1-5 were assigned based on pneumonia severity and IHC staining in the bronchial epithelium and alveoli. Average scores for each group, and the percent of the lung affected by pneumonia, is indicated.

Lastly, because challenge of hamsters with SARS-CoV-2 by IN instillation of liquid viral stocks may underestimate the degree of protective efficacy^41^, we simulated a more physiologic acquisition of SARS-CoV-2 by challenging hamsters were challenged by exposure to an infected hamster on the opposite side of a barrier that permits air exchange but does not permit contact. Naïve hamsters were infected with SARS-CoV-2 Delta (B.1.617.2) two days before being co-housed with two hamsters vaccinated 268 days prior. Vaccinated and naïve-infected hamsters remained on the other side of the barrier for eight hours. To test protection, oropharyngeal swabs and weights were collected up to 14 days post-challenge. When challenged, hamsters immunized with IN Ad4-Wuhan had less weight loss than animals that received Ad4-H5 (p=0.002) or Ad26.CoV2.S (p=0.041)(Figure 7A). Weight loss in Ad26.CoV2.S-immunized animals was not different than animals that received the Ad4-H5 control virus. Animals immunized with IN Ad4-Wuhan had lower total levels of viral replication compared to those immunized with Ad4-H5 (Median AUC: Ad4-Wuhan = 1.5 x 10^5^ vs. Ad4-H5 = 3.8 x 10^7^, p=0.002) or Ad26.CoV2.S (1.13 x 10^7^, p=0.004) (Figure 7B). Animals immunized with IN Ad4-Wuhan also had lower peak viral replication compared to Ad4-H5 or Ad26.CoV.S immunized animals (Max copies/ml: Ad4-Wuhan = 1.6 x 10^5^: Ad4-H5 = 1.3 x 10^7^; Ad26.CoV2.S = 5.4 x 10^6^) (Figure 7C). In one third of Ad4-Wuhan immunized animals (2 of 6) challenge virus replication was not detected.

**Figure 7.**
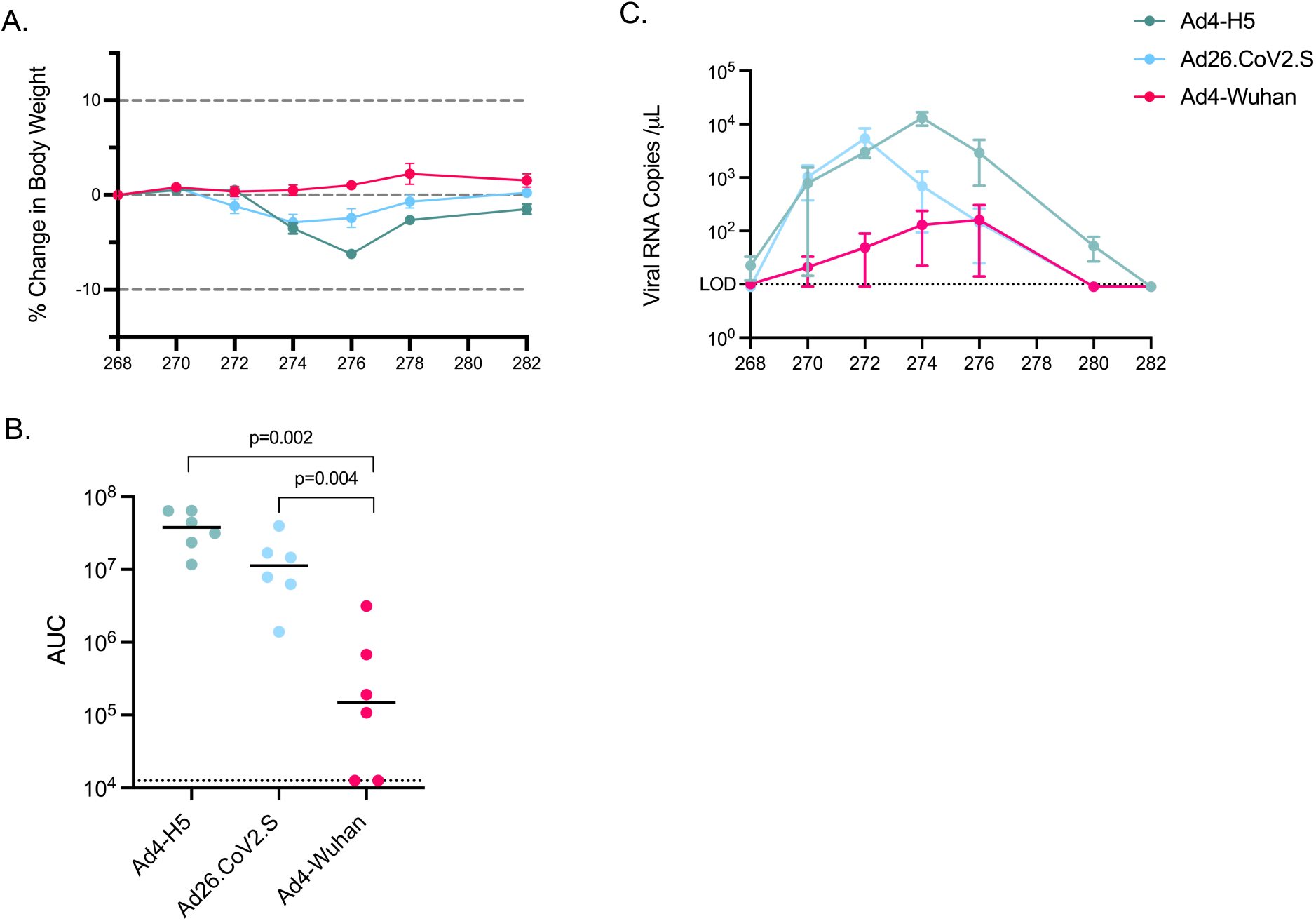
IN Ad4-Spike induces durable restriction of SARS-CoV-2 replication after transmission. Weight loss and viral replication data following transmission at day 268. (A) Average percent body weight relative to the day of challenge, represented as in Fig.4. (B) AUC of viral replication of individual hamsters represented as in Fig.5. Values for each hamster are indicated with dots, the median AUC value for each group is represented as a black line. The dotted line indicates the AUC limit of detection over 14 days (12,600 copies). (C) Curves representing viral replication over time after challenge. The limit of detection is depicted by the dotted line (10 copies per μl). Dots represent medians and bars represent standard error.

## Discussion

This study demonstrates numerous favorable characteristics of the IN Ad4 vector that make it a promising platform for further development as a vaccine for SARS-CoV-2 or other respiratory pathogens with pandemic potential. Hamsters inoculated with Ad4 recombinants replicated the virus and expressed transgenes for at least five days. Although this is below the 7-21 days we have commonly observed in humans, it was sufficient to induce a mucosal immune response, in contrast to IM vaccinations. IN Ad4 also induced a systemic immune response that was similar in magnitude to mRNA-1273 but was considerably more durable. A combined mucosal immune response and a durable systemic response was associated with better outcomes following SARS-CoV-2 challenge in that IN Ad4-S immunized animals consistently had the least weight loss, lowest challenge virus replication, and lowest pathology scores compared to those immunized with IM Ad26.CoV2.S or mRNA1273. This reduced weight loss and lower challenge virus replication was observed after day 174 in an IN challenge study and day 268 in a transmission study demonstrating durable protection. Taken together with clinical data in our prior work and other contexts, these results show that IN Ad4 recombinants have considerable potential to induce durable mucosal and systemic immunity in humans that could provide some protection against disease or infection and diminish the likelihood of transmission if used for SARS-CoV-2, or potentially other respiratory pathogens.

These results add to a growing body of data that show the benefits of respiratory tract delivery of adenoviral recombinant vaccines for SARS-CoV-2. In one recent report, an Ad26-based bivalent (Wuhan and Omicron BA1) vaccine was mucosally administered to Rhesus macaques as a successful boost immunization^13^. Intratracheal instillation of that vaccine provided near-complete protection from a SARS-CoV-2 BQ.1.1 challenge, whereas other boosting strategies, including IN bivalent mRNA were less effective. A single dose of an aerosolized gorilla Ad-S recombinant was also shown to induce protective immunity in Rhesus macaques with impressive durability, dramatically reducing viral load in the bronchoalveolar lavage 64 weeks after immunization^42^. Lastly, when used as a boost following mRNA immunization, a bivalent chimpanzee adenoviral-vectored vaccine (WA1 and BA.5) delivered by an IN mist or inhaled aerosol induced more durable serum antibodies and greater restriction of an Omicron lineage challenge than groups boosted with IM mRNA^15^. However, an intranasally administered chimpanzee adenovirus vectored vaccine induced inconsistent mucosal and modest mucosal and systemic immune responses in one clinical trial, possibly related to the lack of vector replication^43^. In addition, Ad5-S vaccines administered by intranasal spray or inhaled aerosols have shown some promise in clinical trials^20,44^. However, in each of these examples replication incompetent adenoviruses were used. Thus, the magnitude and durability of the immune response likely underestimates those that might be observed in humans following immunization with a replicating vector.

There are reasons to believe that the level of protection observed in the current study may underestimate the protective effect in a clinical setting. First, animals were challenged by directly dripping 10^4^ PFU of SARS-CoV-2 into the nose. This method likely results in a greater delivered dose and higher likelihood of infection and inoculation of the lower respiratory tract compared to contact with aerosols like those used in animal transmission studies here and in prior work. Second, hamsters are only semi-permissive for Ad4 infection. Infection with an Ad4 recombinant lasted less than seven days, much less than the 2-3 weeks observed in seronegative humans^25^. In our experience lower replication is associated with less durable immune responses, such as the in the case of Ad4 recombinants given by the oral route^25^. Even given these caveats, the magnitude of the systemic response to Ad4 in hamsters was equivalent to mRNA at day 56 and higher at day 174. It will be particularly interesting to observe whether administration of Ad4-S to humans provides stable levels of systemic neutralizing antibodies, similar to our experience with Ad4-H5 recombinants.

There is growing evidence that the protective efficacy of mucosal immunity to SARS-CoV-2 is a considerably more formidable barrier to infection and is more durable than previously appreciated. In a study in healthcare workers, prior to the emergence of the genetically distant Omicron variant, all breakthrough infections occurred in mRNA-immunized participants compared to none in those with hybrid immunity from vaccination and prior infection^22^. Further, in a human challenge study SARS-CoV-2 naïve participants 53% had sustained infection following IN inoculum of only 10 50% tissue culture infectious dose (TCID_50_) of a pre-alpha SARS-CoV-2 strain^45^. In contrast, when a similar study was conducted with the same challenge stock in SARS-CoV-2 experienced participants, only 14% of participants had transient infection with challenge doses up to 10^5^ TCID_50_^23^. Mucosal secretory IgA has been found to be the best correlate of protection of the upper respiratory tract against influenza virus, respiratory syncytial virus, and coronaviruses, but levels wane several months after infection. However, clinical studies of IN administration of 10^8^ viral particles of Ad4-H5 to Ad4 seropositive people that are Ad4 seropositive suggest considerable durability based on shortening of adenoviral shedding to less than a week, compared to seronegative participants that shed for 2-3 weeks. This curtailed shedding occurs even though it is likely that their prior infection occurred years earlier. Taken together these data suggest that upper respiratory tract protection or reduction in replication can be durable even though mucosal antibodies have waned. It is likely that reduced replication may be mediated by a rapid recall of T- and B-cell responses.

In the context of prior work with Ad4-H5 in humans, data from the present study suggest that IN replicating Ad4 recombinants represent a promising platform for SARS-CoV-2 and possibly other respiratory pathogens. Of the 32 platforms currently in clinical studies as vaccines for SARS-CoV-2 only an attenuated SARS-CoV-2 (CoviLiv™,Codagenix), a newcastle disease virus vector (NDV-HXP-S, Castlevax), and a parainfluenza virus 5 vector (CVXGA,CyanVac), are replication-competent^17,46-48^. In our experience, and those of others, persistence of antigen that may be provided by replication is an important determinant of immune response durability. These replicating platforms also have extraordinary advantages in manufacturing scalability, and can be self-administered-important characteristics of vaccine platforms in the context of an epidemic or pandemic. It will be important to carefully observe the development of these vaccines for those features of the individual platforms that give the best response regarding protection from infection, lowering transmission, and response durability. The better understanding of these features that comes from comparisons of platforms will likely provide critical lessons for fighting the current and future pandemics.

## Methods

### Construction, production, and purification of Ad4-spike viruses

Six SARS-CoV-2 Spike variant shuttle vector constructs were designed: Wuhan B (GenBank Accession #YP_009724390.1), Beta B.1.351 (GenBank Accession #WZN11859.1 with additional mutations from other circulating Beta strains (L18F, R243I)), Delta B.1.617 (GenBank accession #QVL77043.1 with additional mutations from other circulating Delta strains (R21T, H1071Q)), Gamma P.1 (GenBank Accession #UCU74035.1), Delta B.1.617.2 (GenBank Accession #UAM21534.1 with additional mutations from other circulating Delta strains (R21T, T77K, E154K, Q216H, E482Q, H1099D)), and Omicron BA.2.75.2 (GenBank Accession #UWY19127.1). Additionally, a Wuhan/South Africa recombinant Spike was designed as Wuhan B with the following mutations from South Africa B.1.351: K417N, E484K, N501Y, and D614G. All Spike variants contained two stabilizing substitutions at K986P and V987P. They were synthesized using oligo synthesis via phosphoramidite chemistry (GenScript, Piscataway, NJ, USA) and homologous recombination in a pUC shuttle vector containing the cytomegalovirus (CMV) promoter, tripartite leader sequence (TPL), Kozak sequence, and bovine growth hormone polyadenylation signal (bGH polyA signal) and codon-optimized for human expression. In the Omicron BA.2.75.2 construct, the insert inhibited adenoviral replication which was partially relieved by a truncation of the cytoplasmic tail.

Adenoviral recombinants were produced as previously described^49^ with some modifications. Briefly, the deletion within E3 of Ad4 (GenBank Accession #AY594254) was modified such that only the 12.1 kDa protein, L4 polyA, and E3 polyA were retained. Spike variants were inserted into the Ad4 genome by homologous recombination in electro-competent BJ5183 E. coli (Agilent Technologies, Santa Clara, CA, USA). Candidates were expanded, purified using the ZR BAC DNA Miniprep Kit (Zymo Research, Irvine, CA, USA), and transformed into DH10B ( Thermo Fisher Scientific, Waltham, MA, USA) or NEB 10-beta competent E. coli (New England Biolabs, Ipswich, MA, USA). Subclones were screened by BamHI restriction enzyme digestion and Sanger sequencing (ACGT Inc., Wheeling IL, USA). The positive clones were scaled up using the EndoFree Maxi Kit [12362] (Qiagen, Venlo, Netherlands) for Ad4-Spike (Ad4-S) recombinant assembly.

Linearized Ad4-S DNA plasmid (5.0 μg) and PEIpro transfection reagent (20 μl) (Polyplus, Illkirch, France) were diluted in Opti-MEM reduced serum medium (Thermo Fisher Scientific) (1 mL total) and incubated at room temperature for 10 min. After a 10-minute complexation, the mixture was added dropwise to a 10 cm cell culture treated dish containing ∼2.5 x10^6^ A549 cells (ATCC, Manassas, VA, USA) in Kaighn’s Modification of Ham’s F-12 (F-12K) Medium (ATCC) supplemented with 10% BenchMark™ fetal bovine serum [100-106] (GeminiBio, West Sacramento, CA, USA) and 0.5 mg/mL penicillin-streptomycin-glutamine (Thermo Fisher Scientific) at 37°C with 5% CO_2_. Three days post-transfection, cells were detached using 0.01 M EDTA in PBS and transferred to T300 flasks for Ad4 assembly and incubated at 37°C with 5% CO_2_ for up to 2 weeks. Flasks were monitored for virus-induced cytopathic effect (CPE) and strip-tested regularly with SAS™ Adeno Test (SA Scientific Ltd., San Antonio, TX, USA). Upon Ad4 virus assembly and rescue, the primary viral stocks were harvested, and stored at -80°C as previously described^49^. To confirm the presence of recombinant SARS-CoV-2 spike in the Ad-4 primary viral stocks, DNA was extracted using the MagMAX Viral/Pathogen Kit (Thermo Fisher Scientific), and the region with the Spike transgene was amplified by polymerase chain reaction (PCR) using 10 pmol of the forward primer 5’–AGCTCTTCACTGGGTTTGCGAC–3’ and reverse primer 5’–GAATCCATCTGAAGAGACGAAGGG–3’, Q5 High-Fidelity DNA Polymerase (New England Biolabs) (20 units/mL), Q5 Reaction Buffer (New England Biolabs), High GC Enhancer (New England Biolabs), and 10 mmol dNTPs (New England Biolabs) in a 50 μL reaction with 30 cycles of 95 °C for 15 sec, 56 °C for 15 sec, and 72 °C for 4 minutes and resolved on a 1.2% agarose gel. Constructs with correct band sizes were further confirmed by Sanger sequencing (ACTG Inc.). The primary viral stock was either used directly, or plaque-purified 3 times and concentrated by tangential flow filtration prior to use in the hamster study. The titers of these Ad4-Spike viral stocks were determined using the AdEasy Titering Kit (Agilent Technologies).

Adenovirus serotype 4 expressing luciferase (Ad4-Luc) was constructed by cloning codon optimized luciferase (GenBank Accession #AB762768.1) into the E3 region of the Ad4 genome using the techniques described above. In vitro expression of luciferase was confirmed by infecting A549 cells with Ad4-Luc at an MOI of ∼30. Two days after infection, cells were collected and lysed via incubation in cell culture lysis buffer (Promega, Madison, WI, USA). Bioluminescence of the cell lysate was measured in a black 96 well plate using a PerkinElmer Victor X2 Multilabel Microplate Reader after addition of Luciferase Assay Reagent.

### Characterization of Spike Expression

A549 cells (7 x 10^5^) were seeded in T-25 flasks 24 h before infection with Ad4-S variants at a multiplicity of infection (MOI) of 0.1. For surface staining cells were harvested using 0.01 M EDTA in PBS 48 h after infection and stained with mouse anti- SARS-CoV-2 spike monoclonal antibody (40592-MM117, Sino Biological, Inc., Beijing, China) (50 μL at 1.0 μg/mL final concentration) and then goat anti-mouse IgG PE-conjugated secondary antibody (BioLegend, San Diego, CA, USA) (100 μL at 2.0 μg/mL final concentration) at 37 °C for 30 minutes each. Cells were then stained using LIVE/DEAD Fixable Violet Dead Cell Stain (Thermo Fisher Scientific) at room temperature for 30 minutes (100 μL at a 1:250 final dilution). For intracellular staining, cells were fixed and permeabilized with Cytofix/Cytoperm (Becton Dickinson, Franklin Lakes, NJ, USA) at 4°C for 20 minutes (250 μL) and stained with mouse anti-adenovirus hexon 8C4 monooclonal antibody APC conjugated (Novus Biologicals, LLC., Centennial, CO, USA) at 4 °C for 30 minutes (100 µL at 1.11 µg/mL final concentration). The following antibodies were used to stain uninfected cells for appropriate compensation controls for each fluorchrome: anti-human CD24-PE (BioLegend) (50 μL at 20 μg/mL final concentration) at 37 °C for 30 min, APC anti-human CD24 (BioLegend) (50 μL at 10 μg/mL final concentration) at 37 °C for 30 min, and LIVE/DEAD Fixable Violet Dead Cell Stain at room temperature for 30 min (100 μL at 1:250 final dilution) were used for compensation controls. Samples were then analyzed by flow cytometry on an BD FACSymphony™S6 Cell Sorter Flow Cytometer (Becton Dickinson) to verify the presence of adenovirus-infected cells that express SARS-CoV-2 Spike.

### Hamster Bioluminescence Assay, Immunization, and Challenge

All animal experiments were conducted according to the animal study proposal (ASP LIR22/LIRID10, LIR27/LIRID11E, LID44E) approved by National Institutes of Health, National Institute of Allergy and Infectious Diseases, Animal Care and Use Committee that meets all federal requirements, as defined in the Animal Welfare Act, NIH Guide for Care and Use of Laboratory Animals (The Guide), the Public Health Service Policy (PHS), and the Humane Care and Use of Laboratory Animals in AALAC accredited facilities. This study used 6-8 week old male Syrian Hamsters [HsdHan®: Aura) (Inotiv, West Lafayette, IN, USA) that were co-housed, all of which were processed for terminal bleed collections under general anesthesia and euthanized by exsanguination as approved by the AVMA (American Veterinary Medical Association) and adopted by NIH-NIAID ACUC.

Bioluminescence was measured from the nose of anesthetized male hamsters (Age ∼18 weeks) via IVIS Spectrum In Vivo Imaging System (Caliper Life Sciences, Hopkinton, MA, USA) after intranasal inoculation with 50 μl of 1.4 x 10^8^ IFU/mL of Ad4 Luc on Day 0. Each hamster was injected (intraperitoneal) with 18mg of D-Luciferin Potassium Salt Bioluminescent Substrate (IVISbrite D-Luciferin Potassium Salt Bioluminescent Substrate (Revvity Health Sciences, Waltham, MA, USA) in normal saline approximately 10 mins prior to IVIS imaging. Average Radiance [photons/sec/cm^2^/steradian] was calculated using the Living Image Software version 4.3.1 (Revvity Health Sciences), after subtracting average radiance of the nose of an uninfected male hamster.

A dose titration of Ad4-Wuhan was conducted in male, 6-8 week old, Syrian hamsters (HsdHan®: Aura) (Inotiv, West Lafayette, IN, USA) by IN immunization with 10^2^-10^7^ IFU dripped onto the nose (25µl per nostril) in six groups (n=6 each). Hamster body weights and oropharyngeal swabs were collected on 0, 1, 3, 5, and 7 days post-immunization. Oropharyngeal swabs were collected using sterile Microbrush Test Swabs (Young Innovations, Algonquin IL, USA) that were placed in Viral Transport Medium (Wuxi NEST Biotechnology Co., Ltd, Jiangsu, China) and stored at -80°C. In subsequent experiments, hamsters (n=6 per group) were immunized IN with 10^7^ IFU of Ad4 recombinants expressing SARS-CoV-2 Wuhan (B), Delta (B.1.617.2), or Omicron (BA.2.75.2) S protein, or Influenza H5 (A/Vietnam/1194/2004) as a negative control. As a comparator, separate groups of hamsters (n=6 per group) were immunized intramuscularly (IM) with two doses of the Moderna mRNA-1273 (5 µg)^50^ on days 0 and 28, or Johnson & Johnson Ad26.CoV2.S vaccine (5 x 10^9^ VP)^51^ in a single dose on day 0. Hamster weights and oropharyngeal swabs were collected every other day for 12-14 days.

To assess the immunogenicity of the Ad4-S recombinants, blood was collected at 0, 28, and 56 DPI from retro-orbital venous plexus, while nasal wash and BAL were collected at the time of euthanasia at day 56. To assess protective efficacy, hamsters immunized with Ad4-S recombinants (n=14 per group) were IN challenged with either SARS-CoV-2 Delta (B.1.617.2) or Omicron (B.1.1.529) (10^4^ PFU) at 56 or 172 days by dripping 25µl of the challenge material in each nostril in an BSL3 laboratory. Challenge viruses were propagated using VeroE6 cells that constitutively express a human transmembrane protease serine 2 (TMPRSS2)^47^ under geneticin selection, at an MOI of 0.01 for 72 hours and validated for sequence and titer as previously described^52^. Hamsters were bled immediately prior to challenge, and hamster weights and oropharyngeal swabs were collected post-challenge every other day for 12-14 days. Four hamsters from each group were sacrificed 4 days after challenge for bronchoalveolar lavage (BAL), nasal wash, and pulmonary histopathology and immunohistochemical analyses.

To assess protection from airborne transmission, hamsters were immunized with IN Ad4-H5 and Ad4-Delta or IM for Ad26.CoV2.S (n=6 per group) as described above. To generate infected hamsters for the airborne transmission study, nine naive hamsters were infected with 10^4^ PFU SARS-CoV-2 Delta (B.1.617.2) by dripping 25 µl in each nostril and allowing 2 days for viral replication. One SARS-CoV-2-infected hamster was placed on one side of a cage and 2 immunized hamsters on the other for 8 hours separated by a barrier that permitted air exchange but no direct contact. Weights and oropharyngeal swabs were collected every other day for 14 days post-challenge. All vaccinations and sample collections were either performed under isoflurane sedation or following euthanasia.

### Pseudovirus Production and Neutralization Assay

Serum neutralizing antibody titers were calculated using a lentivirus-based assay as a function of the reduction in luciferase gene expression following a single round of infection with SARS-CoV-2 spike-pseudotyped viruses in an HEK293T/ACE2 expressing cell line (obtained from Dr. Michael Farzan, The Scripps Research Institute), as described^53^. Briefly, SARS-CoV-2 pseudoviruses were produced by transfection of HEK-293T/17 (ATCC, Manassas, VA, USA) cells with plasmids encoding a lentivirus backbone vector (pCMV ΔR8.2), firefly luciferase reporter (pHR’ CMV Luc)^54^, TMPRSS2^55^, and a full-length SARS-CoV-2 spike from Wuhan (B.1.1), Delta (B.1.617.2), Omicron (B.1.1.529) or Omicron (BA.2.75.2) using FuGENE 6 transfection reagent (Promega, Madison, WI, USA). Plasmids encoding SARS-CoV-2 spike proteins were obtained from Dr. Nicole Doria-Rose at the NIH Vaccine Research Center. Pseudoviruses were titered to determine the the dilution producing 200,000 relative luminescence units (RLU) that was then used in the assay. Pseudoviruses were co-incubated with 8 serial 5-fold dilutions of heat-inactivated serum samples (1:20 or 1:30 starting dilution) in cell culture-treated 96-well plates for 45 min at 37°C, prior to the addition of HEK293T/ACE2 cells (10^4^ cells/well). Following incubation for 72 h at 37°C, cells were disrupted on a platform shaker at 600 rpm in Luciferase cell culture lysis reagent (Promega). Luciferase was then activated with Bright-Glo Luciferase Assay System (Promega) and the RLU was measured on a Spectramax L luminometer (Molecular Devices, San Jose, CA, USA). Serum neutralization titers were calculated as the inhibitory dilution of serum at which RLU are determined to be 50% (ID_50_) compared to virus-only control wells after subtracting cell only background luminescence^53^. The limit of detection was an ID_50_ of 30. Sera with no detectable neutralization above background were assigned an ID_50_ of 29.

### Viral Nucleic Acid Quantification

Quantification of Ad4-S during the immunization phase and SARS-CoV-2 virus during the challenge phase was determined by quantitative PCR (qPCR). Ad4-S viral DNA was extracted using the MagMAX DNA Multi-Sample Ultra Kit (Thermo Fisher Scientific) according to the manufacturer’s protocol. Briefly, 50 µL of the viral transport media used to elute oropharyngeal swabs was mixed with 450 µL of DNA Lysis Buffer and 300 µL of 100% isopropanol by vortexing, and 40 µL of DNA magnetic binding beads were added. Beads containing bound DNA were RnaseA treated and then washed before DNA elution. qPCR reactions were prepared using TaqMan Fast Advanced Master Mix (Thermo Fisher Scientific) with the forward primer 5’-GTCTCTGCTGATCGTGAACAA-3’, reverse primer 5’-GGACTCCATCCAGCTCTTATTG-3’, and probe 5’ (6-FAM)-TCATCAAGGTGTGCGAGTTCCAGT-(TAMRA-Sp) 3’. The reactions conditions were as follows: 95°C for 20 s, followed by 40 cycles of 1 s at 95°C and 20 s at 60°C. Amplifications were performed using a QuantStudio 3 Real Time PCR instrument and analyzed with QuantStudio 3/5 Real-Time PCR Software through ThermoFisher Connect Platform (Thermo Fisher Scientific).

SARS-CoV-2 RNA was extracted with the Direct-zol-96 MagBead RNA extraction kit (Zymo Research, Irvine CA, USA) according to the manufacturer’s protocol. To extract SARS-CoV-2 RNA from challenged hamsters, 50 µL of viral transport media from oropharyngeal swabs was mixed with 150 µL of Trizol-LS (Thermo Fisher Scientific) for inactivation of SARS-CoV-2. The remainder of the RNA extraction was performed using a KingFisher Flex System (Thermo Fisher Scientific). In 96-well round deep well plates, 200 µL of sample was combined with 200 µL ethanol before adding 20 µL MagBinding Beads to capture viral RNA. Beads were Dnase I treated and washed sequentially with MagBead DNA/RNA Wash buffer 1 and 2 before eluting viral RNA. qPCR reactions were performed with TaqMan Fast Virus 1-Step Master Mix (Thermo Fisher Scientific), forward primer (5’-CGATCTCTTGTAGATCTGTTCTC-3’) in the 5’ leader region and gene-specific probes (5’ (6-FAM)-CGATCAAAACAACGTCGGCCCC-(BHQ1) 3’) and reverse primer (5’-GGTGAACCAAGACGCAGTAT-3’) as, previously described^56^. The reaction conditions were as follows: 50°C for 5 min, 95°C for 20 s, and 40 cycles of 95°C for 15 s and 60°C for 1 min. Amplifications were performed with a QuantStudio 6 Pro Real-Time PCR System (Thermo Fisher Scientific). The assay limit of detection was 5 copies per reaction or 1000 copies per ml of viral transport medium, determined by serially diluted subgenomic SARS-CoV-2 RNA transcript spiked into medium, similarly extracted and amplified.

### Multiplexed Ig Binding Assay

Binding antibodies to SARS-CoV-2 S antigen in hamster serum, nasal wash, and BAL were quantified using a custom multiplex bead array, as previously described^57^. Briefly, SARS-CoV-2 S antigens, including RBD: Alpha (B1.1.7), Beta (B.1.351) and Omicron (B.1.1.529); S2: (Alpha (B1.1.7) and Beta (B1.351); and S: Alpha (B.1.1.7), Delta (B.1.617.2), Gamma (P.1), Omicron (B.1.1.529), were covalently coupled with Luminex MagPlex® Magnetic Microspheres (Diasorin, Saluggia, Italy) for the detection of Spike-binding antibodies. SARS-CoV-1 S and SARS-CoV2 nucleoprotein were included as non-specific antigen controls. Antibody isotypes were detected using rabbit anti-hamster IgG, IgA, or IgM secondary antibodies conjugated to R-phycoerythrin (Brookwood Biomedical, Jemison, AL, USA). Serum and nasal wash dilutions were optimized in pilot experiments, with test concentrations of serum ranging from 1:250 to 1:5000, and concentrations of nasal wash and BAL ranging from 1:10 to 1:100. Mean Fluorescence Intensity was measured on a FlexMap 3D® or xMAP Intelliflex® Systems (Diasorin).

### Lung Histopathology and Immunohistochemistry

For histopathology, lung tissues were fixed in formalin before being dehydrated with graded alcohols, cleared in xylene, and infiltrated with paraffin. Tissues were then embedded in paraffin and cut on a microtome at 5 µm. For immunohistochemistry, slides were deparaffinized in xylene and hydrated with graded alcohols. Slides were then subject to antigen retrieval in a sodium citrate buffer (pH 6) and blocked against endogenous peroxidases. Following blocking, slides were incubated with the primary antibody (GenScript, U86YFA140-4/CB2093). Slides were then stained with a goat-anti-rabbit horseradish peroxidase secondary (Vector Laboratories, MP-6401) and visualized with diaminobenzidine. Finally, slides were counterstained with hematoxylin, dehydrated with graded alcohols, cleared in xylene, and mounted with permanent mounting media. All washes were performed using either distilled water or TBST at room temperature.

Slides were scored in a blinded manner. Percent Lung Affected was estimated to 5% of the total tissue area using Spot Imaging Solutions version 5.2.7. Pneumonia severity and IHC staining was scored from 0-5, with 0 representing normal lungs. Scores between 1 and 5 represent tissues with minimal (1-3) foci of inflammation to severe inflammation with most lobes impacted. IHC scores in the alveoli and bronchial epitheliuim represent lungs with rare/small foci of immunoreactivity to tissues where almost all airways are affected with significant immunoreactivity.

### Statistics

The Wilcoxon rank sum tests were used for pairwise comparisons in Fig 2, 3, 5, 6, 7A. For comparisons of body weight changes over time, comparisons were done of the area under the curve for the weight change trajectory during the period of interest for Fig 1A, 4, 7A, and Supp. Fig 1. No adjustment for multiple comparisons was performed. The analyses were performed using custom scripts in R (v 4.5.0).

## Data Availability

Data is provided within the supplementary information files.

## Acknowledgements

This research was supported by the Intramural Research Program of the National Institute of Allergy and Infectious Diseases. It also has been funded in part with federal funds from the National Cancer Institute, National Institutes of Health, under Contract No. 75N91019D00024. The content of this publication does not necessarily reflect the views or policies of the Department of Health and Human Services, nor does mention of trade names, commercial products, or organizations imply endorsement by the U.S. Government. We would like to thank the entire staff of the NIH Division of Veterinary Resources and the entire animal care staff for their support of these experiments.

## Author Contributions

K.M., R.F.J., P.F.W., M.E.A., D.D., J.I.C., and M.C. contributed to the conception and design of the project; M.C. and R.J. supervised the work. A.P., F.V., R.R., E.K., T.S., J.X., and P.M.R. contributed to the construction of vaccine candidates. A.C.V. and A.H. performed *in* situ immunofluorescence imaging to determine Ad4 replication. J.A.W., G.K., S.C.S., T.K., and I.S. developed and performed the microplex bead assay for binding antibody detection. B.K., I.S, M.H., and E.O. optimized and performed serum neutralization assays. K.M., E.W., T.B., S.G.C., R.F.J., I.S., and P.M.R. participated in conduct of animal experiments. B.K., I.S., E.O., R.R, P.M.R., C.C., J.D.W., and T.M. processed oropharyngeal swabs, and M.G., Z.Z., K.R., M.R.B, and E.M. performed qPCR quantification of viral RNA. V.C. and J.L. performed statistical analysis. B.K, I.S., R.R., P.M.R, T.K, and M.C. assisted with preparation of the manuscript.

